# Inference, validation and predictions about statistics and propagation of cortical spiking in vivo

**DOI:** 10.1101/363085

**Authors:** J. Wilting, V. Priesemann

## Abstract

Electrophysiological recordings of spiking activity can only access a small fraction of all neurons simultaneously. This spatial subsampling has hindered characterizing even most basic properties of collective spiking in cortex. In particular, two contradictory hypotheses prevailed for over a decade: the first proposed an asynchronous irregular, the second a critical state. While distinguishing them is straightforward in models, we show that in experiments classical approaches fail to infer them correctly, because subsampling can bias measures as basic as the correlation strength. Deploying a novel, subsampling-invariant estimator, we find evidence that in vivo cortical dynamics clearly differs from asynchronous or critical dynamics, and instead occupies a narrow “reverberating” regime, consistently across multiple mammalian species and cortical areas. These results enabled us to predict cortical properties that are difficult or impossible to obtain experimentally, including responses to minimal perturbations, intrinsic network timescales, and the strength of external input compared to recurrent activation.

## Introduction

When investigating spiking activity in neuronal networks, only a tiny fraction of all neurons can be recorded experimentally with millisecond precision. Such spatial subsampling fundamentally limits virtually any recording and hinders inferences about the collective dynamics of cortical networks.^1–4^ In fact, even some of the most basic characteristics of cortical network dynamics are not known with certainty, such as the population Fano factor, or the fraction of spikes generated internally versus those triggered by input.

In particular, two contradicting hypotheses to describe network dynamics have competed for more than a decade, and are the subjects of ongoing scientific debate: One hypothesis suggests that collective dynamics are “asynchronous irregular”^5–7^ (AI), i.e. neurons spike independently of each other and in a Poisson manner, which may reflect a balanced state.^8,9^ The other hypothesis proposes that neuronal networks operate at criticality^10–17^ and thus in a particularly sensitive state close to a phase transition. These hypotheses have distinct implications for the coding strategy of the brain. The typical balanced state minimizes redundancy,^18–22^ supports fast network responses,^8^ and shows vanishing autocorrelation time (*τ* → 0). In contrast, criticality in models optimizes performance in tasks that profit from extended reverberations of activity in the network,^23–29^ because it is characterized by long-range correlations in space and time (*τ* → ∞). It has been proposed that *τ* reflects an integration window over past activity, thereby allowing brain networks to operate on specific timescales.^30–33^ Timescales estimated from single neurons span hundreds of milliseconds,^34^ but it is unclear how timescales of the full network can be inferred in the face of subsampling.

Surprisingly, there is experimental evidence for both AI and critical states in cortical networks, although both states are clearly distinct. Evidence for the AI state is based on characteristics of single neuron spiking resembling a Poisson process, i.e. exponential inter spike interval (ISI) distributionss and Fano factors *F* close to unity.^35^ Moreover, spike count crosscorrelations^36,37^ are small. Evidence for criticality was typically obtained from a population perspective instead, and assessed neuronal avalanches, i.e. spatio-temporal clusters of activity,^1,10,38–41^ whose sizes are expected to be power-law distributed if networks are critical.^42^ Deviations from power-laws, typically observed for spiking activity in awake animals,^2,3,43,44^ were attributed to subsampling effects.^1–4,45–47^ Hence, different analysis approaches provided evidence for one or the other dynamical state’s dominance.

We rely on a classic approach to probe the dynamical states of a system at steady state, namely applying minimal perturbations. Studying how perturbations cascade through a system enables the inference of numerous system properties. London and colleagues applied such a perturbation framework and estimated that one average *m* = 28 additional postsynaptic spikes are triggered by one extra spike in a presynaptic neuron from intracellular recordings.^48^ From their complementary extracellular spike recordings, one can equally well estimate *m* ≈ 0.04 Hz/neuron · 10 ms · *k* = 0.6: in the 10 ms subsequent to the perturbation, an increase of 0.04 Hz is observed for each neuron. Assuming that this 10 ms is an upper bound to directly activate any of the *k* ≈ 1500 directly connected neurons, one obtains as as upper bound *m* ≈ 0.04 Hz/neuron · 10 ms · *k* = 0.6. This vast range for estimates of *m* arises largely because such inferences are heavily influenced by subsampling. We here build on a subsampling-invariant approach presented in a companion study,^49^ which allows us to resolve questions surrounding the contradictory results on cortical dynamics: (i) we establish an analytically tractable minimal model for *in vivo* like activity, which can interpolate from AI to critical dynamics; (ii) we estimate the dynamical state of cortical activity based on a novel, subsampling-invariant estimator;^49^ (iii) we predict a number of network dynamical properties, which are experimentally accessible and allow to validate our approach; (iv) we predict a number of yet unknown network properties, including *m*, the expected number of spikes triggered by one additional spike, the emergent network timescale *τ*, the distribution of the total number of spikes triggered by a single extra action potential, and the fraction of activation that can be attributed to afferent external input to a cortical network.

## Material and Methods

### Minimal model of spike propagation

To gain an intuitive understanding of our mathematical approach, make a thought experiment in your favorite spiking network: apply one additional spike to an excitatory neuron, in analogy to the approach by London and colleagues^48^. How does the network respond to that perturbation? As a first order approximation, one quantifies the number of spikes that are triggered by this perturbation *additionally* in all postsynaptic neurons. This number may vary from trial to trial, depending on the membrane potential of the postsynaptic neurons; however, what interests us most is *m*, the *mean number of spikes triggered by the one extra spike*. Taking a mean-field approximation and assuming that the perturbation indeed is small, any of these triggered spikes in turn trigger spikes in their postsynaptic neurons in a similar manner, and thereby the perturbation may cascade through the system. Mathematically, such cascades can be described by a branching process.^50–52^

In the next step, assume that perturbations are started continuously at rate *h*, for example through afferent input from other brain areas or sensory modalities. As neurons presumably do not distinguish whether a postsynaptic potential was elicited from a neuron from within the network, or from afferent input, all spikes are assumed to have on average the same impact on the network dynamics. Together, this leads to the mathematical framework of a branching network,^2,3,10,24,45^ which can generate dynamics spanning AI and critical states depending on the input,^53^ and hence is well suited to probe network dynamics *in vivo* (see Supp. 1 for details). Most importantly, this framework allows to infer *m* and other properties from the ongoing activity proper, because one treats any single spike as a minimal perturbation on the background activity of the network. Mathematical approaches to infer *m* are long known if the full network is sampled.^54,55^ Under subsampling, however, it is the novel estimator described in^49^ that for the first time allows an unbiased inference of *m*, even if only a tiny fraction of neurons is sampled. After inferring *m*, a number of quantities can be analytically derived, and others can be obtained by simulating a mean-field spiking model, which is constrained by the experimentally measured *m* and the spike rate.

The framework of branching networks can be interpreted as a stochastic description of spike propagation on networks, as outlined above. It can alternatively be taken as a *strictly phenomenological* approximation to network dynamics that enables us to infer details of network *statistics* despite subsampling. Independent of the perspective, the dynamics of the network is mainly governed by *m* (Fig. 1a). If an action potential only rarely brings any postsynaptic neuron above threshold (*m* ⪆ 0), external perturbations quickly die out, and neurons spike independently and irregularly, driven by external fluctuations *h*. In general, if one action potential causes less than one subsequent action potential on average (*m* < 1), perturbations die out and the network converges to a stable distribution, with increasing fluctuations and variance the closer *m* is to unity. If *m* > 1, perturbations may grow infinitely, potentially leading to instability. The critical state (*m* = 1) separates the stable (subcritical) from the unstable (supercritical) phase. When approaching this critical state from below, the expected size 〈*s*〉 and duration 〈*d*〉 of individual cascades or avalanches diverge: 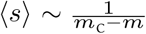. Therefore, especially close to criticality, a correct estimate of *m* is vital to assess the risk that the network develops large, potentially devastating cascades, which have been linked to epileptic seizures,^56^ either generically or via a minor increase in *m*.

**Figure 1:**
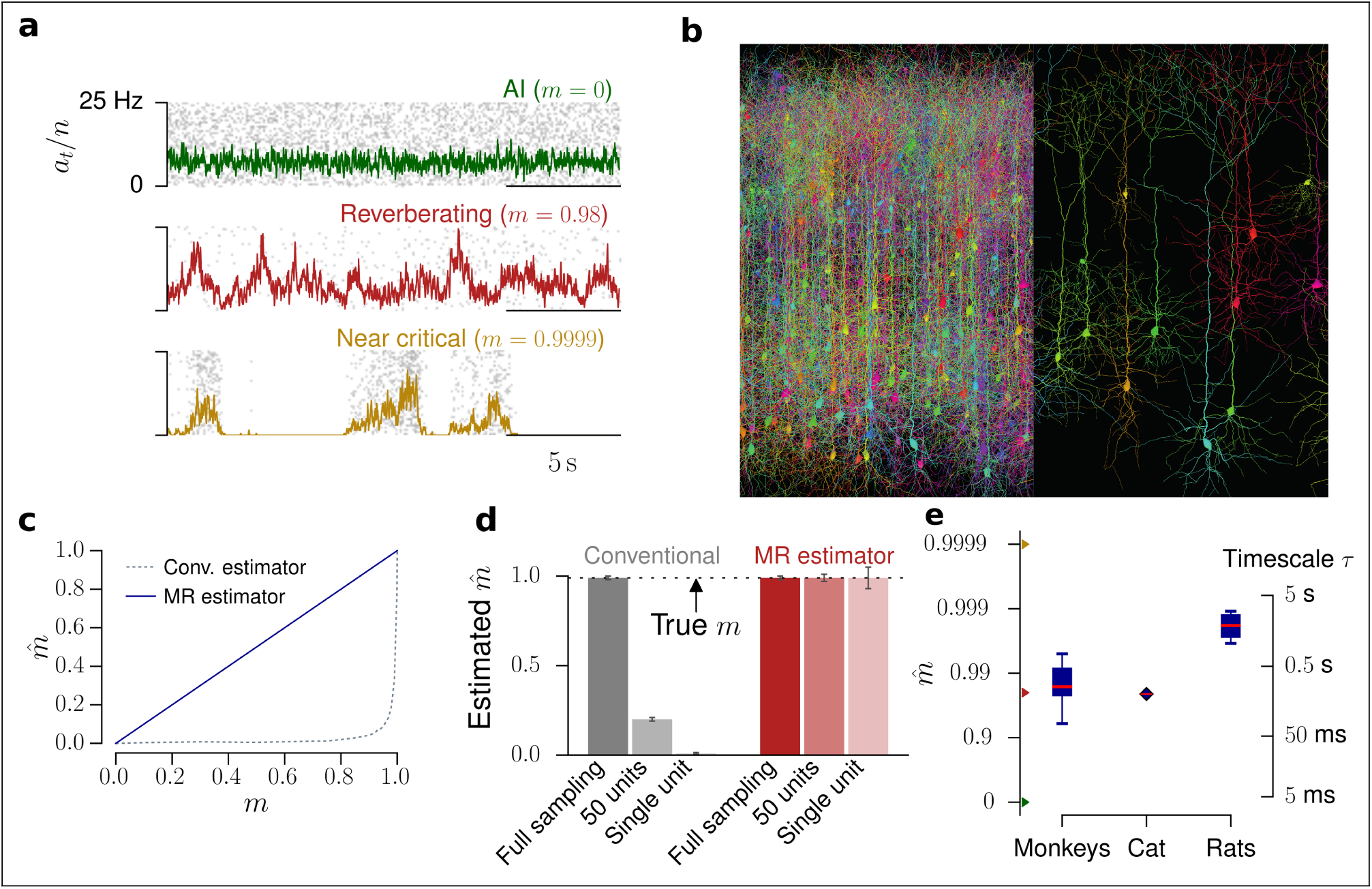
Spatial subsampling. **a**. Raster plot and population rate for networks with different spike propagation parameters. They exhibit vastly different dynamics, which readily manifest in the population activity. **b**. When assessing neuronal spiking activity, only a small subset of all neurons can be recorded. This spatial subsampling can hinder correct inference of collective properties of the whole network; figure created using TREES^65^ and reproduced from^49^. **c**. Estimated branching ratio *m*̂ as a function of the simulated branching ratio *m*, inferred from subsampled activity (100 out of 10,000 neurons). While the conventional estimator misclassified *m* from this subsampled observation (gray, dotted line), the novel multistep regression (MR) estimator returned the correct values **d**. For a reverberating branching network with *m* = 0.98, the conventional estimator inferred *m*̂ = 0.21 or *m*̂ = 0.002 when sampling 50 or 1 units respectively, in contrast to MR estimation, which returned the correct *m*̂ even under strong subsampling. **e**. Using the novel MR estimator, cortical network dynamics in monkey prefrontal cortex, cat visual cortex, and rat hippocampus were consistently found to exhibit reverberating dynamics, with 0.94 < *m*̂ < 0.991 (median *m*̂ = 0.98 over all experimental sessions, boxplots indicate median / 25% – 75% / 0% – 100% over experimental sessions per species). These correspond to network timescales between 80 ms and 2 s.

#### Simulation

We simulated a branching network model by mapping a branching process^50^ (Supp. 1) onto a fully connected network of *N* = 10, 000 neurons.^24^ An active neuron activated each of its *k* postsynaptic neurons with probability *p* = *m*/*k*. Here, the activated postsynaptic neurons were drawn randomly without replacement at each step, thereby avoiding that two different active neurons would both activate the same target neuron. Similar to the branching process, the branching network is critical for *m* = 1 in the infinite size limit, and subcritical (supercritical) for *m* < 1 (*m* > 1). We modeled input to the network at rate h by Poisson activation of each neuron at rate *h*/*N*. Subsampling^1^ was applied to the model by sampling the activity of *n* neurons only, which were selected randomly before the simulation, and neglecting the activity of all other neurons.

If not stated otherwise, simulations were run for *L* = 10^7^ time steps (corresponding to ~11 h). Confidence intervals were estimated according to^49^ from *B* = 100 realizations of the network, both for simulation and experiments.

The reverberating branching networks were defined to match the respective experimental recording in the number of sampled neurons *n*, mean activity 〈*a*_*t*_〉, and branching ratio *m*. Exemplarily for the cat recording, which happened to represent the median *m*̂, this yielded *m* = *m*̂ = 0.98, *n* = 50, and 〈*a*_*t*_〉 = 1.58 per bin, from which *h* = 0.032 · *N* follows. The corresponding AI and near-critical networks were matched in *n* and 〈*a*_*t*_〉, but set up with branching ratios of *m* = 0 or *m* = 0.9999 respectively. For all networks, we chose a full network size of *N* = 10^4^.

In Figs. 2c, the reverberating branching network was also matched to the length of the cat recording of 295 s. To test for stationarity, the cat recordings and the reverberating branching network were split into 59 windows of 5 s each, before estimating *m* for each window. In Fig. 1c, subcritical and critical branching networks with *N* = 10^4^ and 〈*A*_*t*_〉 = 100 were simulated, and *n* = 100 units sampled.

**Figure 2:**
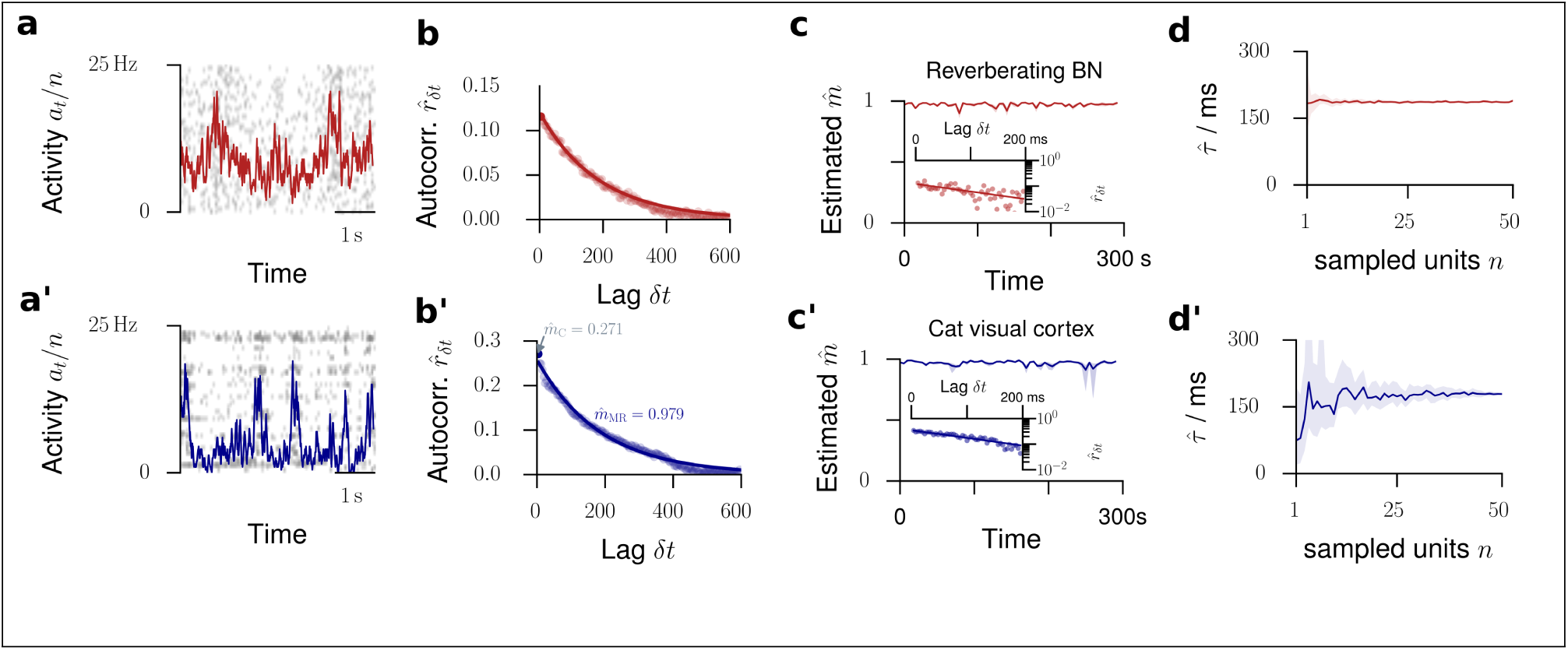
Validation of the model assumptions. The top row displays properties from a reverberating network model, the bottom row spike recordings from cat visual cortex. **a/a’**. Raster plot and population activity *a*_*t*_ within bins of Δ*t* = 4 ms, sampled from *n* = 50 neurons. **b/b’**. Multistep regression (MR) estimation from the subsampled activity (5 min recording). The predicted exponential relation *r*_*δt*_ ~ *m*^*δt*/Δ*t*^ = exp(–*δt*/*τ*) provides a validation of the applicability of the model. The experimental data are fitted by this exponential with remarkable precision. **c/c’**. The estimated branching parameter *m*̂ for 59 windows of 5 s length suggests stationarity of *m* over the entire recording (shaded area: 16% to 84% confidence intervals). The variability in *m*̂ over consecutive windows was comparable for experimental recording and the matched network (*p* = 0.09, Levene test). Insets: MR estimation exemplified for one example window each. **d/d’**. When subsampling even further, MR estimation always returns the correct timescale *τ*̂ (or *m*̂) in the model. In the experiment, this invariance to subsampling also holds, down to *n* ≈ 10 neurons (shaded area: 16% to 84% confidence intervals estimated from 50 subsets of *n* neurons).

### Experiments

We evaluated spike population dynamics from recordings in rats, cats and monkeys. The rat experimental protocols were approved by the Institutional Animal Care and Use Committee of Rutgers University.^57,58^ The cat experiments were performed in accordance with guidelines established by the Canadian Council for Animal Care.^59^ The monkey experiments were performed according to the German Law for the Protection of Experimental Animals, and were approved by the Regierungspräsidium Darmstadt. The procedures also conformed to the regulations issued by the NIH and the Society for Neuroscience. The spike recordings from the rats and the cats were obtained from the NSF-founded CRCNS data sharing website.^57–60^

#### Rat experiments

In rats the spikes were recorded in CA1 of the right dorsal hippocampus during an open field task. We used the first two data sets of each recording group (ec013.527, ec013.528, ec014.277, ec014.333, ec015.041, ec015.047, ec016.397, ec016.430). The data-sets provided sorted spikes from 4 shanks (ec013) or 8 shanks (ec014, ec015, ec016), with 31 (ec013), 64 (ec014, ec015) or 55 (ec016) channels. We used both, spikes of single and multi units, because knowledge about the identity and the precise number of neurons is not required for the MR estimator. More details on the experimental procedure and the data-sets proper can be found in^57,58^.

#### Cat experiments

Spikes in cat visual cortex were recorded by Tim Blanche in the laboratory of Nicholas Swindale, University of British Columbia.^59^ We used the data set pvc3, i.e. recordings of 50 sorted single units in area 18.^60^ We used that part of the experiment in which no stimuli were presented, i.e., the spikes reflected spontaneous activity in the visual cortex of the anesthetized cat. Because of potential non-stationarities at the beginning and end of the recording, we omitted data before 25 s and after 320 s of recording. Details on the experimental procedures and the data proper can be found in^59,60^.

#### Monkey experiments

The monkey data are the same as in^3,61^. In these experiments, spikes were recorded simultaneously from up to 16 single-ended micro-electrodes (ⵁ = 80*μ*m) or tetrodes (ⵁ = 96*μ*m) in lateral prefrontal cortex of three trained macaque monkeys (M1: 6 kg ♀ M2: 12 kg ♂; M3: 8 kg ♀). The electrodes had impedances between 0.2 and 1.2 M*Ω* at 1 kHz, and were arranged in a square grid with inter electrode distances of either 0.5 or 1.0 mm. The monkeys performed a visual short term memory task. The task and the experimental procedure is detailed in^61^. We analyzed spike data from 12 experimental sessions comprising almost 12.000 trials (M1: 5 sessions; M2: 4 sessions; M3: 3 sessions). 6 out of 12 sessions were recorded with tetrodes. Spike sorting on the tetrode data was performed using a Bayesian optimal template matching approach as described in^62^ using the “Spyke Viewer” software.^63^ On the single electrode data, spikes were sorted with a multi-dimensional PCA method (Smart Spike Sorter by Nan-Hui Chen).

### Analysis

#### Temporal binning

For each recording, we collapsed the spike times of all recorded neurons into one single train of population spike counts *a*_*t*_, where *a*_*t*_ denotes how many neurons spiked in the *t*^*th*^ time bin *Δt*. If not indicated otherwise, we used *Δt* = 4 ms, reflecting the propagation time of spikes from one neuron to the next.

#### Multistep regression estimation of *m*̂

From these time series, we estimated *m*̂ using the MR estimator described in^49^. For *k* = 1,…, *k*_max_, we calculated the linear regression slope *r*_*kΔt*_ for the linear statistical dependence of *a*_*t*+*k*_ upon *a*_*t*_. From these slopes, we estimated *m*̂ following the relation *r*_*δt*_ = *b* · *m*̂^*δt*/Δ*t*^, where *b* is an (unknown) parameter that depends on the higher moments of the underlying process and the degree of subsampling. However, for an estimation of *m* no further knowledge about *b* is required.

Throughout this study we chose *k*_max_ = 2500 (corresponding to 10 s) for the rat recordings, *k*_max_ = 150 (600 ms) for the cat recording, and *k*_max_ = 500 (2000 ms) for the monkey recordings, assuring that *k*_max_ was always in the order of multiple auto-correlation times.

In order to test for the applicability of a MR estimation, we used a set of conservative tests^49^ and included only those time series, where the approximation by a branching network was considered appropriate. For example, we excluded all recordings that showed an offset in the slopes *r*_*k*_, because this offset is, strictly speaking, not explained by a branching network and might indicate non-stationarities. Details on these tests are found in^49^. Even with these conservative tests, we found the exponential relation *r*_*k*_ = *bm*^*δt*/Δ*t*^ expected for branching networks in the majority of experimental spike recordings (14 out of 21, Fig. S1).

#### Avalanche size distributions

Avalanche sizes were determined similarly to the procedure described in^1,3^. Assuming that individual avalanches are separated in time, let {*t*_*i*_} indicate bins without activity, *a*_*t*_*i*__ = 0. The size *s*_*i*_ of one avalanche is defined by the integrated activity between two subsequent bins with zero activity:

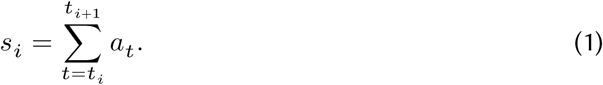

From the sample {*s*_*i*_} of avalanche sizes, avalanche size distributions *p*(*s*) were determined using frequency counts. For illustration, we applied logarithmic binning, i.e. exponentially increasing bin widths for *s*.

For each experiments, these empirical avalanche size distributions were compared to avalanche size distributions obtained in a similar fashion from three different matched models (see below for details). Model likelihoods *l*({*s*_*i*_})|*m*) for all three models were calculated following^64^, and we considered the likelihood ratio to determine the most likely model based on the observed data.

#### ISI distributions, Fano factors and spike count cross-correlations

For each experiment and corresponding reverberating branching network (subsampled to a single unit), ISI distributions were estimated by frequency counts of the differences between subsequent spike times for each channel.

We calculated the single unit Fano factor *F* = Var[*a*_*t*_]/〈*a*_*t*_〉 for the binned activity *a*_*t*_ of each single unit, with the bin sizes indicated in the respective figures. Likewise, single unit Fano factors for the reverberating branching networks were calculated from the subsampled and binned time series.

From the binned single unit activities 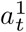 and 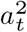 of two units, we estimated the spike count cross correlation 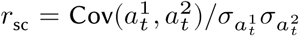. The two samples 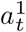 and 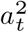 for the reverberating branching networks were obtained by sampling two randomly chosen neurons.

## Results

### Subsampling-invariant inference of the dynamical state

In a companion study^49^ we showed that conventional estimators based on linear regression^54,55^ significantly underestimate *m*̂ when the system is subjected to subsampling (Fig. 1c), as it is always the case in electrophysiological recordings (Fig. 1b). The bias is considerable: For example, sampling 50 neurons or a single neuron in a branching network with *m* = 0.99 resulted in the wrong estimates *m*̂_Conv_ = 0.21, or even *m*̂_Conv_ = 0.002, respectively (Fig. 1d). Thus a process close to instability (*m* = 0.99) is mistaken as Poisson-like (*m*̂_Conv_ = 0.002 ≈ 0) just because the estimate from subsampled activity is taken as face value for the entire population. The same study presented a novel *multistep regression* estimator (MR estimator), which correctly characterizes the population dynamics via *m* even under strong subsampling, in principle even from single neurons. Importantly, one can estimate *m* even when sampling only a very small fraction of neurons and without knowing the network size *N*, the number of sampled neurons *n*, nor any moments of the underlying process.^49^ This robustness makes the estimator an ideal tool for the analysis of neuronal network recordings.

### Reverberating spiking activity *in vivo*

We analyzed *in vivo* spiking activity from Macaque monkey prefrontal cortex during a short term memory task,^61^ anesthetized cat visual cortex with no stimulus,^59^ and rat hippocampus during a foraging task.^57,58^ We applied MR estimation to the binned population spike counts *a*_*t*_ of the recorded neurons of each session (see methods). In the continuous spectrum from AI (*m* = 0) to critical (*m* = 1), we identified a limited range of branching values *in vivo*: in the experiments *m*̂ ranged from 0.963 to 0.998 (median *m*̂ = 0.98), corresponding to autocorrelation times between 100 ms and 2 s (median 247 ms, Figs. 1e, S1). This clearly suggests that spiking activity *in vivo* is neither AI-like, nor consistent with a critical state. Instead, it is poised in a regime that, unlike critical or AI, does not maximize one particular property alone but may combine features of both (see discussion). Due to the lack of one prominent characterizing feature, we name it the *reverberating* regime, stressing that activity reverberates (different from the AI state) at timescales of hundreds of milliseconds (different from a critical state, where they can persist infinitely).

### The reverberating state differs from criticality

On first sight, *m*̂ = 0.98 of the reverberating state may suggest that the collective spiking dynamics is very close to critical. Indeed, physiologically a *Δm* ≈ 1.6% difference to criticality is small in terms of the effective synaptic strength. However, this apparently small difference in single unit properties has a large impact on the collective *dynamical* fingerprint and makes AI, reverberating, and critical states clearly distinct: (1) This distinction is readily manifest in the fluctuations of the population activity, where states with *m* = 0.98 and *m* = 0.999 are clearly different (Fig. 1a). (2) Consider the sensitivity to a small input, i.e. the susceptibility 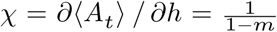. The susceptibility diverges at criticality. A critical network is thus overly sensitive to input. In contrast, states with *m* ≈ 0.98 assure sensitivity without instability. (3) Likewise, the *Δm* ≈ 1.6% difference limits the intrinsic timescale of the dynamics to a few hundred milliseconds, while at criticality it approaches infinity. (4) Because of the divergences at criticality, network dynamics dramatically differ between *m* = 0.9, *m* = 0.99 or *m* = 0.999: for example, the differences in susceptibility (sensitivity) and variance are 100-fold. Because this has a strong impact on network dynamics and putative network function, finely distinguishing between dynamical states is both important and feasible even if the corresponding differences in effective synaptic strength (*m*) appear small.

### Validity of the approach

There is a straight-forward verification of the validity of our phenomenological model: it predicts an exponential autocorrelation function *r*_*δt*_ for the population activity *a*_*t*_. We found that the activity in cat visual cortex (Figs. 2a,a’) is surprisingly well described by this exponential fit (Fig. 2b,b’). This validation holds to the majority of experiments investigated (14 out of 21, Fig. S1).

A second verification of our approach is based on its expected invariance under subsampling: We further subsampled the activity in cat visual cortex by only taking into account spikes recorded at a subset of *n* out of all available single units. As predicted (Fig. 2d), the estimates of *m*̂, or equivalently of *τ*̂, coincided for any subset of single units. Only if the activity of less than 5 out of the available 50 single units was considered, the autocorrelation time was underestimated (Fig. 2d’), most likely because of the heterogeneity of cortical networks. These results demonstrate, however, that our approach gives consistent results from the activity of *n* ≥ 5 neurons, which were available for all investigated experiments.

### Origin of the activity fluctuations

The fluctuations found in cortical spiking activity, instead of being intrinsically generated, could in principle arise from non-stationarities, which could in turn lead to misestimation of *m*. This is unlikely for three reasons: First, we defined a set of conservative tests to reject recordings that show any signature of common non-stationarities. Even with these tests, we found the exponential relation *r*_*δt*_ ~ *m*^*δt*/Δ*t*^ expected for branching networks not only in cat visual cortex, but in the majority of experiments (14 out of 21, Fig. S1). Second, recordings in cat visual cortex were acquired in absence of any stimulation, excluding stimulus-related non-stationarities. Third, when splitting the spike recording into short windows, the window-to-window variation of *m*̂ in the recording did not differ from that of stationary *in vivo*-like branching networks (*p* = 0.3, **Figs. 2c,c’**). The *in vivo*-like branching network by definition was set up with the same branching ratio *m*, spike rate 〈*a*_*t*_〉, number of sampled neurons *n*, and duration as the experimental recording (e.g. for the cat *n* = 50, *m* = 0.98, *r*̄ = 7.9 Hz, recording of 295 s length). For these reasons the observed fluctuations likely reflect intrinsic timescales of the underlying collective network dynamics.

### Timescales of the network and single units

The dynamical state described by *m* directly relates to an exponential autocorrelation function with an intrinsic network timescale of *τ* = –*Δt*/ ln *m*. Exemplarily for the cat recording, *m* = 0.98 implies a network timescale of *τ* = 188 ms, where we here chose *Δt* = 4 ms. While the autocorrelation function of the full network activity is expected to show an exponential decay, we showed that the autocorrelation of single neuron activity rapidly decreases at the timescale of a bin size (Fig. 3a). This rapid decrease is typically interpreted a lack of memory, overlooking that single neurons do not need to be equivalent to the network in terms of autocorrelation strength. Our theoretical results explain how this prominent dip comes about even in reverberating systems: because of the strong subsampling when considering single neuron activity, the strength of autocorrelation is decreased by a constant factor for any lag *δt* ≠ 0. Ignoring the value at *δt* = 0, the floor of the autocorrelation function still unveils the exponential relation. Remarkably, the autocorrelogram of single units in cat visual cortex displayed precisely the shape of autocorrelation predicted for single neurons (compare Figs. 3a and b).

**Figure 3:**
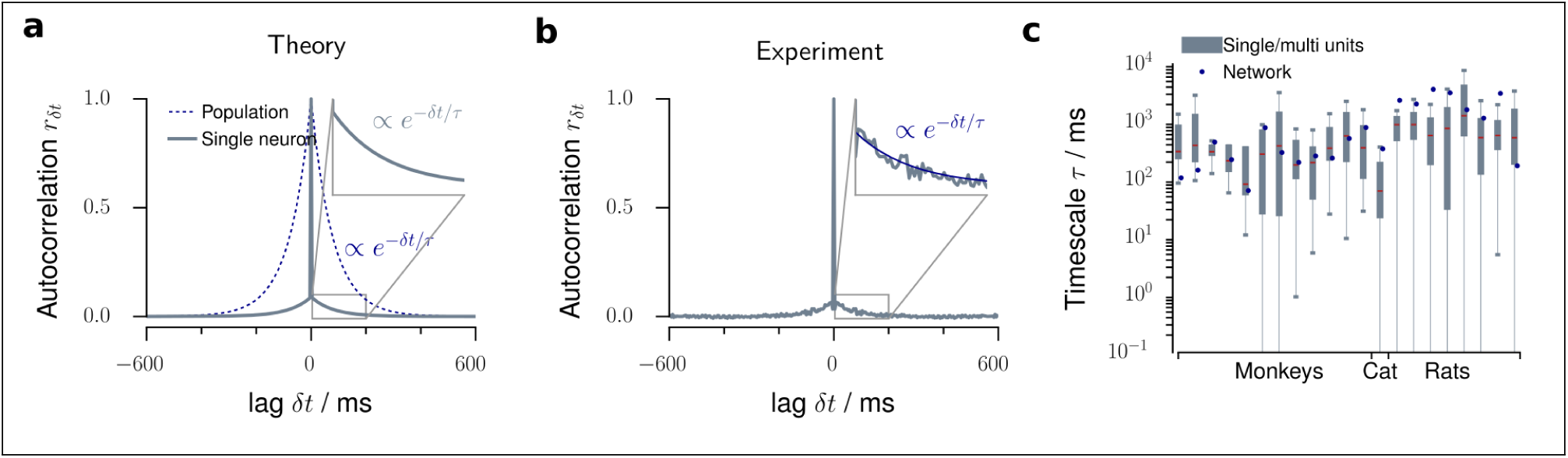
MR estimation and autocorrelation times. **a**. In a branching model, the autocorrelogram of the population activity is exponential with decay time *τ* (blue dotted line). In contrast, the autocorrelogram for single neurons shows a sharp drop from *r*_0_ = 1 at lag *δt* = 0s to the next lag *r*_±*Δt*_ (gray solid line). We showed that this drop is a subsampling-induced bias. When ignoring the zero-lag value, the autocorrelation strength is decreased, but the exponential decay and even the value of the autocorrelation time *τ* of the network activity are preserved (inset). **b**. The autocorrelogram of single neuron activity recorded in cat visual cortex precisely resembles this theoretical prediction, namely a sharp drop and then an exponential decay. **c**. Single unit and population timescales for all experimental sessions. The boxplots indicate the distribution of timescales inferred from single unit activity (median in red / 25% – 75% / 2.5% – 97.5%), the blue dots the timescale inferred from the population activity of all sampled units.

Although our results were largely invariant to further subsampling (provided *n* ≥ 5, Fig. 2d’), the intrinsic timescales *τ* (or *m*) of single neurons differed from the network timescale, as one might expect in heterogeneous systems. We found that single neuron timescales were typically smaller than the network timescale (Fig. 3c, median *τ* = 85 ms for single neurons in cat visual cortex versus *τ* = 180 ms for the network, Figs. 2d’, S9c). Therefore, the network timescale inferred by our approach contributes further information about network dynamics compared to previous studies which only considered single neurons.^34^

### Established methods are biased to identifying AI dynamics

On the population level, networks with different *m* are clearly distinguishable (Fig. 1a). Surprisingly, single neuron statistics, namely interspike interval (ISI) distributions, Fano factors, conventional estimation of *m*, and the autocorrelation *r*_*δt*_, all returned signatures of AI activity regardless of the underlying network dynamics and cannot serve as a reliable indicator for the network’s dynamical state.

First, exponential interspike interval (ISI) distributions are considered a strong indicator of Poisson-like firing. Surprisingly, the ISIs of single neurons in the *in vivo*-like branching network closely followed exponential distributions, which were determined mainly by the firing rate, and were almost indistinguishable from ISI distributions obtained from AI networks (Figs. 4a,a’, S2). This result was confirmed by coefficients of variation close to unity, as expected for exponential distributions (Fig. S2).

**Figure 4:**
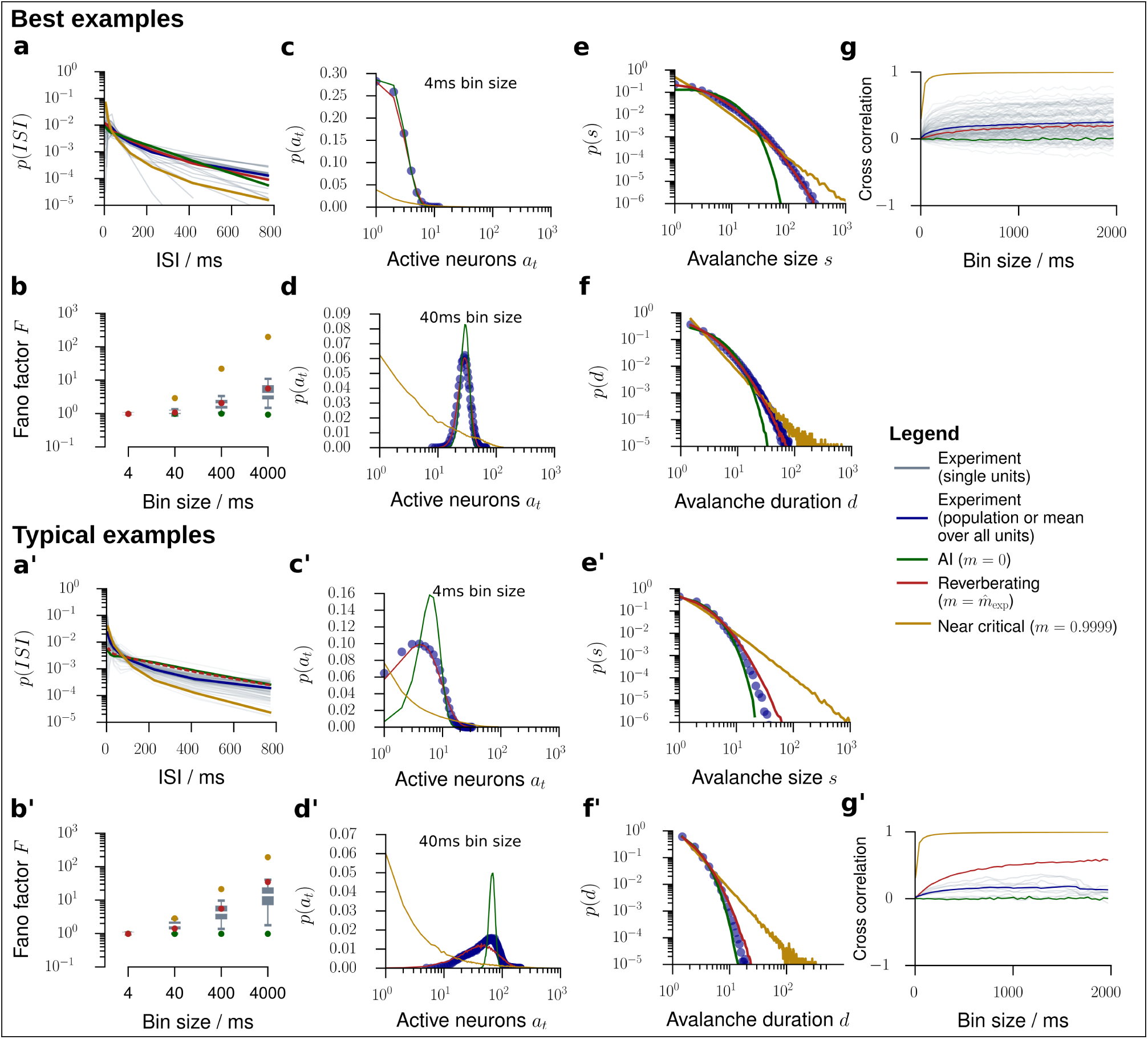
Model validation for *in vivo* spiking activity. We validated our model by comparing experimental results to predictions obtained from the *in vivo*-like, reverberating model, which was matched to the recording in the mean rate, inferred *m*, and number of recorded neurons. In general, the experimental results (gray or blue) were best matched by this reverberating model (red), compared to asynchronous irregular (AI, green) and critical (yellow) models. From all experimental sessions, best examples (top) and typical examples (bottom) are displayed. For results from all experimental sessions see Figs. S2 to S8. **a/a’**. Inter-spike-interval (ISI) distributions. **b/b’**. Fano factors of single neurons for bin sizes between 4 ms and 4s. **c/c’**. Distribution of spikes per bin *p*(*a*_*t*_) at a bin size of 4 ms. **d/d’**. Same as **c/c’** with a bin size of 40 ms. **e/e’**. *In vivo* avalanche size distributions *p*(*s*) for all sampled units. AI activity lacks large avalanches, near critical activity produces power-law distributed avalanches, even under subsampling. **f/f’**. *In vivo* avalanche duration distributions *p*(*d*) for all sampled units. textbfg/**g’**. Spike count cross-correlations (*r*_sc_) as a function of the bin size.

Second, the Fano factor *F* for the activity of single neurons was close to unity, a hallmark feature of irregular spiking,^35^ in any network model (Fig. 5g, analytical result: Eq. (S8)) and for single unit activity across all units and experiments (Figs. 4b,b’, S3). Even when increasing the bin size to 4s, the median Fano factor of single unit activity did not exceed *F* = 10 in any of the experiments, even in those with the longest reverberation. In contrast, for the full network the Fano factor rose to *F* ≈ 10^4^ for the *in vivo*-like branching network and diverged when approaching criticality (Fig. 5g, analytical result: Eq. (S4)).

**Figure 5:**
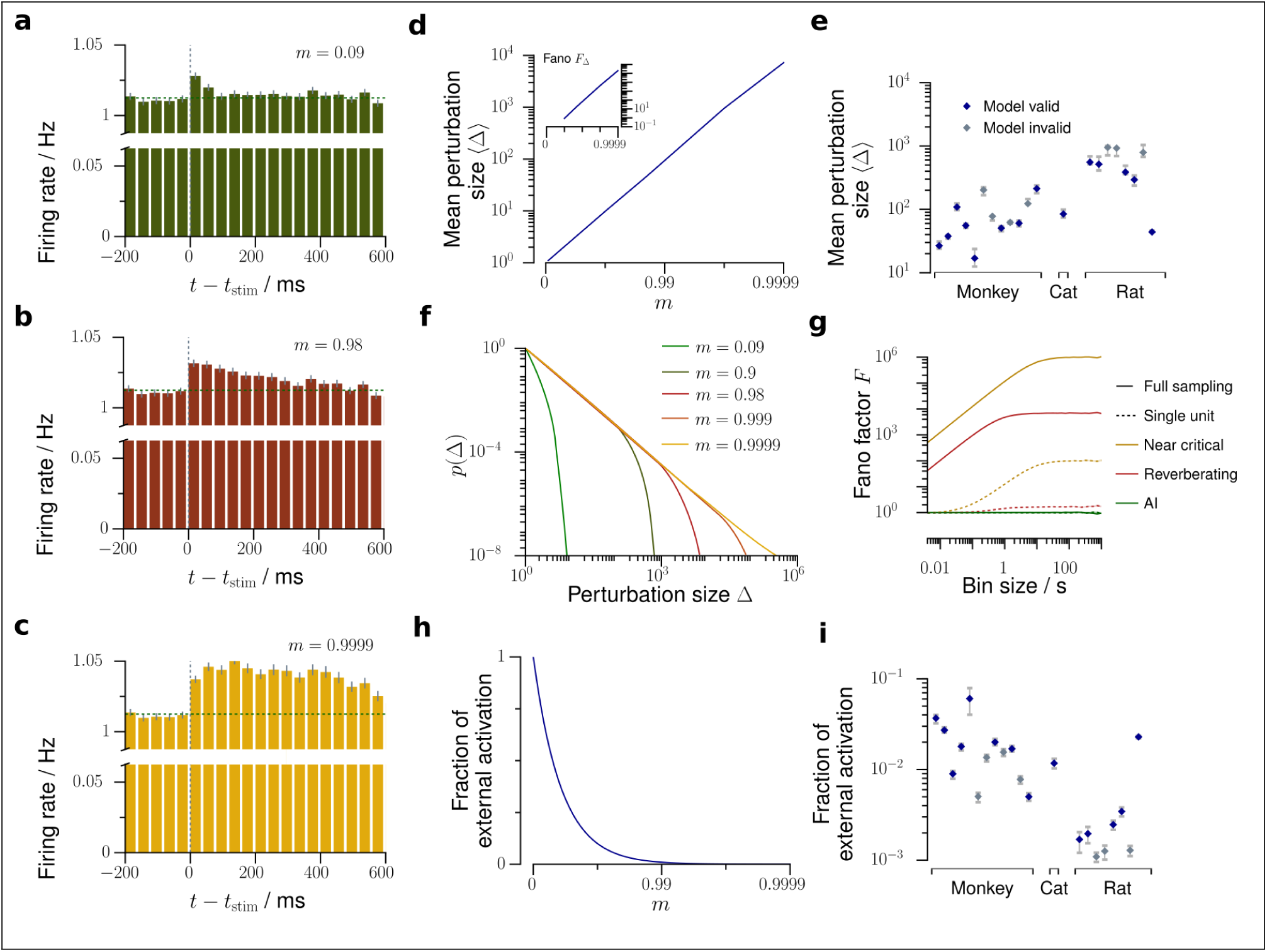
Predictions about network activity. Using our *in vivo*-like, reverberating model, we can predict several network properties, which are yet very hard or impossible to obtain experimentally. **a - c**. In response to one single extra spike, a perturbation propagates in the network depending on the branching ratio *m* and can be observed as a small increase of the average firing rate of the sampled neurons, here simulated for 500 trials (see also London et al. [48]). This increase of firing rate decays exponentially, with the decay time *τ* being determined by *m*. The perturbation **a** is rapidly quenched in the asynchronous irregular network, **b** decays slowly over hundreds of milliseconds in the reverberating state, or **c** persists almost infinitely in the critical state. **d**. The average perturbation size 〈*Δ*〉 and Fano factor *F*_*Δ*_(inset) increase strongly with *m*. **e**. Average total perturbation sizes predicted for each spike recording of mammalian cortex (errorbars: 5% – 95% confidence intervals). **f**. Distribution *p*(*Δ*) of total perturbation sizes *Δ*. The asynchronous irregular networks show approximately Poisson distributed, near critical networks power-law distributed perturbation sizes. textbfg. Bin size dependent Fano factors of the activity, here exemplarily shown for the asynchronous irregular (*m* = 0, green), representative reverberating (*m* = 0.98, red), and near critical (*m* = 0.9999, yellow) networks. While the directly measurable Fano factor of single neurons (dotted lines) underestimates the Fano factor of the whole network, the model allows to predict the Fano factor of the whole network (solid lines). **h**. The fraction of the externally generated spikes compared to all spikes in the network strongly decreases with larger *m*. **i**. Fraction of the externally generated spikes predicted for each spike recording of mammalian cortex (errorbars as in **e**).

Third, conventional regression estimators^54,55^ are biased towards inferring irregular activity, as shown before. Here, conventional estimation yielded a median of *m*̂ = 0.057 for single neuron activity in cat visual cortex, in contrast to *m*̂ = 0.954 returned by MR estimation even from single unit recordings (Fig. S9).

Fourth, when examining the autocorrelation function of an experimental recording (Fig. 3b) the prominent decay of *r*_*τt*_ prevails and hence single neuron activity appears uncorrelated in time.

### Cross-validation of model predictions

We compared the experimental results to an *in vivo*-like model, which was matched to the recording only in the average firing rate of single neurons, and in the inferred branching ratio m. Remarkably, this *in vivo*-like branching network could predict statistical properties not only of single neurons as shown before (ISI and Fano factor, see above), but also pairwise and population properties. This prediction capability further underlines the usefulness of this simple model to approximate the ground state of cortical dynamics.

First, the model predicted the activity distributions, *p*(*a*_*t*_), better than AI or critical networks for the majority of experiments (15 out 21, Figs. 4c,d,c’,d’, S5, S6), both for the exemplary bin sizes of 4 ms and 40 ms. Hence, branching networks only matched in their respective first moments of the activity distributions (through the rate) and first moments of the spreading behavior (through *m*) in fact approximated all higher moments of the activity distributions *p*(*a*_*t*_).

Likewise, the model predicted the distributions of neural avalanches, i.e. spatio-temporal clusters of activity (Figs. 4e,f,e’, f’, S7, S8). Characterizing these distributions is a classic approach to assess criticality in neuroscience,^1,10^ because avalanche size and duration distributions, *p*(*s*) and *p*(*d*) respectively, follow power laws in critical systems (yellow). In contrast, for AI activity, they are approximately exponential^66^ (green). The matched branching networks predicted neither exponential nor power law distributions for the avalanches, but very well matched the distributions of the experiment (compare red and blue). Indeed, model likelihood^64^ favored the *in vivo*-like branching network over Poisson and critical networks for the majority experiments (18 out of 21, Fig. S7). Our results here are consistent with those of spiking activity in awake animals, which typically do not display power laws.^2,3,43^ In contrast, most evidence for criticality has been based on *coarse* measures of neural activity (LFP, EEG, BOLD; see^3^ and references therein).

Last, the model predicted the pairwise spike count cross correlation *r*_sc_. In experiments, *r*_sc_ is typically between 0.01 and 0.25, depending on brain area, task, and most importantly, the analysis timescale (bin size).^37^ For the cat recording the model even correctly predicted the bin size dependence of *r*_sc_ from *r*̄_sc_ ≈ 0.004 at a bin size of 4 ms (analytical result: Eq. (S11)) to *r*̄_sc_ ≈ 0.3 at a bin size of 2 s (Fig. 4g). Comparable results were also obtained for some monkey experiments. In contrast, correlations in most monkey experiments and rat hippocampal neurons showed smaller correlation than predicted (Figs. 4g’, S4). It is very surprising that the model correctly predicted the cross-correlation even in some experiments, as *m* was inferred only from the *temporal* structure of the spiking activity alone, whereas *r*_sc_ characterizes spatial dependencies.

Overall, by only estimating the effective synaptic strength *m* from the *in vivo* recordings, higher-order properties like avalanche size distributions, activity distributions and in some cases spike count cross correlations could be closely matched using the generic branching network.

### The dynamical state determines responses to small stimuli

After validating the model using a set of statistical properties that are well accessible experimentally, we now turn to making predictions for yet unknown properties, namely network responses to small stimuli. In the line of London and colleagues^48^, assume that on a background of spiking activity one single extra spike is triggered. This spike may in turn trigger new spikes, leading to a cascade of additional spikes *Δ*_*t*_ propagating through the network. A dynamical state with branching ratio *m* implies that *on average*, this perturbation decays with time constant *τ* = –*Δt*/log*m*. Similar to the approach in^48^, the evolution of the mean firing rate, averaged over a reasonable number of trials (here: 500) unveils the nature of the underlying spike propagation: depending on *m*, the rate excursions will last longer, the higher *m* (Figs. 5a,b,c, S10a). The perturbations are not deterministic, but show trial-to-trial variability which also depends on *m* (S10b).

Unless *m* > 1, the theory of branching networks ensures that perturbations will die out eventually after a duration *d*, having accumulated a total of 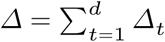 extra spikes in total. This perturbation size *Δ* and duration *d* follow specific distributions,^50^ which are determined by *m*: they are power law distributed in the critical state, with a cutoff for any *m* < 1 (Fig. 5f, Supplementary Figs. S10c,d). These distributions imply a characteristic perturbation size 〈*Δ*〉 (Fig. 5d), which diverges at the critical point. The variability of the perturbation sizes is also determined by *m* and also diverges at the critical point (inset of Fig. 5d and Supplementary Fig. S10e).

Taken together, these results imply that the closer a neuronal network is to criticality, the more sensitive it is to external perturbations and can better amplify small stimuli. At the same time, these networks also show larger trial-to-trial variability. For typical cortical networks, we found that the response to one single extra spike will on average comprise between 20 and 1000 additional spikes in total (Figs. 5e).

### The dynamical state determines network susceptibility and variability

Moving beyond single spike perturbations, our model gives precise predictions for the network response to continuous stimuli. If extra action potentials are triggered at rate *h* in the network, the network will again amplify these external activations, depending on *m*. Provided an appropriate stimulation protocol, this rate response could be measured and our prediction tested in experiments (Fig. S10g). The susceptibility d*r*/d*h* diverges at the critical transition and is unique to a specific branching ratio *m*. We predict that typical cortical networks will amplify a small, but continuous increase of the input rate about 50-fold (Fig. S10h, red).

While the mean activity is determined by the network input and its susceptibility, the network activity fluctuates around this mean value. The magnitude of these fluctuations in relation to the mean can be described by the network Fano factor *F* = Var[*A*_*t*_]/〈*A*_*t*_〉 (Fig. 5g). This quantity cannot be directly inferred from experimental recordings, because the Fano factor of subsampled populations severely underestimates the network Fano factor, as shown before. We here used our *in vivo*-like model to obtain estimates of the network Fano factor: for a bin size of 4 ms it is about *F* ≈ 40 and rises to *F* ≈ 4000 for bin sizes of several seconds.

### Distinguishing afferent and recurrent activation

Last, our model gives an easily accessible approach to solving the following question: given a spiking neuronal network, which fraction of the activity is generated by recurrent activation from within the network, and which fraction can be attributed to external, afferent excitation? The branching model readily provides an answer: the fraction of externally generated activity is *h*/〈*A*〉 = 1 – *m* (Fig. 5h). In this framework, AI-like networks are completely driven by external input currents or noise, while reverberating networks generate a substantial fraction of their activity intrinsically. For the experiments investigated in this study, we inferred that between 0. 1% and 7% of the activity are externally generated (median 2%, Fig. 5i). While this view may be simplistic given the complexity of neuronal network activity, keep in mind that “all models are wrong, but some are useful”.^67^ Here, the model has proven to provide a good first order approximation and therefore promises to make reasonable predictions on properties of spiking networks.

## Discussion

### Our results resolve contradictions between AI and critical states

Our results for spiking activity *in vivo* suggest that network dynamics shows AI-like statistics, because under subsampling the observed correlations are underestimated. In contrast, typical experiments assessing criticality potentially overestimated correlations by sampling from overlapping populations (LFP, EEG) and thereby hampered a fine distinction between critical and subcritical states.^68^ By employing for the first time a consistent, quantitative estimation, we provided evidence that *in vivo* spiking population dynamics reflects a reverberating state, i.e. it lives in a narrow regime around *m* = 0.98. This result is supported by the findings by Dahmen and colleagues:^69^ based on distributions of covariances, they inferred that cortical networks should operate in a regime below criticality. Given the generality of our results across different species, brain areas, and cognitive states, our results suggest self-organization to this regime as a general organization principle for neural network dynamics.

### The reverberating state combines features of AI and critical regimes

Operating in a reverberating state, which is between AI and critical, may combine the computational advantages of the two dynamical states: (1) AI networks react to external input rapidly, and show very little reverberation of the input. In contrast, criticality is associated with “critical slowing down”, i.e. performing any computation might take overly long. The *m* = 0.98 state shows intermediate timescales of a few hundred milliseconds. These reverberations may carry short term memory and allow to integrate information over limited timescales.^27,70^ (2) Criticality has been associated with maximal processing capacity. However, a number of everyday tasks, e.g. memory recall, require only sufficient capacity for survival and reproduction rather than maximum capacity.^71^ Thus maximizing this one property alone is most likely not necessary from an evolutionary point of view. One particular example manifest from our results is the trade-off between sensitivity and reliability: while the critical state maximizes sensitivity by amplifying small stimuli (Fig. 5h), this sensitivity comes at the cost of increased trial-to-trial variability (Fig. 5i) and therefore may hinder reliable responses.^72^ (3) Criticality in a branching process marks the transition to unstable dynamics. These instabilities have been associated with epilepsy.^56^ The prevalence of epilepsy in humans^73,74^ supports our results that the brain indeed operates biophysically still close to instability, but keeps a sufficient safety-margin to make seizures sufficiently unlikely.^3^ This is in line with our results that the effective synaptic strength is close to, but not at *m* = 1.

### More complex network models

Cortical dynamics is clearly more complicated than a simple branching network. For example, heterogeneity of neuronal morphology and function, non-trivial network topology, and the complexity of neurons themselves are likely to have a profound impact on the population dynamics. However, we showed that *statistics* of cortical networks are well approximated by a branching network. Therefore, we interpret branching networks as a *statistical* approximation of spike propagation, which can capture dynamics as complex as cortical activity. By using branching networks, we draw on the powerful advantage of analytical tractability, which allowed for basic insight into dynamics and stability of cortical networks.

It is a logical next step to refine the model by including additional relevant parameters, guided by the results obtained from the well-understood estimator. For example, our results show that networks with balanced excitation and inhibition,^8,75,76^ which became a standard model of neuronal networks,^77^ should be extended to incorporate the network reverberations observed *in vivo*. Possible candidate mechanisms are increased coupling strength or inhomogeneous connectivity. Both have already been shown to exhibit rate fluctuations with timescales of several hundred milliseconds.^78–80^

Likewise, neuron models of spike responses typically model normally distributed network synaptic currents, which originate from the assumption of uncorrelated Poisson inputs. Our results suggest that this input should rather exhibit reverberating properties with timescales of a few hundred milliseconds to reflect input from cortical neurons *in vivo*.

### Deducing network properties from the tractable model

Using the tractable model, we could predict and validate network properties, such as distributions of avalanche sizes and durations, interspike intervals, or activities. Given the experimental agreement with these predictions, we deduced further properties, which are impossible or difficult to assess experimentally and gave insight into more complex questions about network responses: how do perturbations propagate within the network and how susceptible is the network to external stimulation?

One particular question we could address is the following: which fraction of network activity is attributed to external or recurrent, internal activation? We inferred that about 98% of the activity are generated by recurrent excitation. However, note that this result likely depends on the brain area and cognitive state investigated: For layer 4 of primary visual cortex in awake mice, Reinhold and colleagues^81^ concluded that the fraction of recurrent cortical excitation rises to only about 72% and cortical activity dies out with a timescale of about 12 ms after thalamic silencing. Their numbers agree perfectly well with our phenomenological model: a timescale of 12 ms implies that the fraction of recurrent cortical activation is *m* ≈ 0.71, just as found experimentally. Under anesthesia, in contrast, they report timescales of several hundred milliseconds, in agreement with our results. These differences show that the fraction of external activation may strongly depend on cortical area, layer, and cognitive state. The novel estimator can in future contribute to a deeper insight into these differences, because it allows for a straight-forward assessment of afferent versus recurrent activation without the requirement of thalamic or cortical silencing.

## Acknowledgments

JW received support from the Gertrud-Reemstma-Stiftung. VP received financial support from the German Ministry for Education and Research (BMBF) via the Bernstein Center for Computational Neuroscience (BCCN) Göttingen under Grant No. 01GQ1005B, and by the German-Israel-Foundation (GIF) under grant number G-2391-421.13. JW and VP received financial support from the Max Planck Society.

## Competing interests

The authors declare that the research was conducted in the absence of any commercial or financial relationships that could be construed as a potential conflict of interest.

## Supplementary material for “Inference, validation and predictions about statistics and propagation of cortical spiking in vivo” by J. Wilting and V. Priesemann

### Supp. 1 Branching processes

In a branching process (BP) with immigration^50–52^ each unit *i* produces a random number *y*_*t*, *i*_ of units in the subsequent time step. Additionally, in each time step a random number *h*_*t*_ of units immigrates into the system (drive). Mathematically, BPs are defined as follows:^50,51^ Let *y*_*t*, *i*_ be independently and identically distributed non-negative integer-valued random variables following a law *Y* with mean *m* = 〈*Y*〉 and variance *σ*^2^ = Var[*Y*]. Further, *Y* shall be non-trivial, meaning it satisfies P[*Y* = 0] > 0 and P[*Y* = 0] + P[*Y* = 1] < 1. Likewise, let h_t_ be independently and identically distributed non-negative integer-valued random variables following a law *H* with mean rate *h* = 〈*H*〉 and variance *ξ*^2^ = Var[*H*]. Then the evolution of the BP *A*_*t*_ is given recursively by

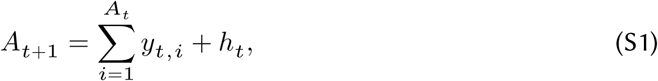

i. e. the number of units in the next generation is given by the offspring of all present units and those that were introduced to the system from outside.

The stability of BPs is solely governed by the mean offspring *m*. In the subcritical state, *m* < 1, the population converges to a stationary distribution *A*_*∝*_ with mean 〈*A*_*∝*_〉 = *h*/(1 – *m*). At criticality (*m* = 1), *A*_*t*_ asymptotically exhibits linear growth, while in the supercritical state (*m* > 1) it grows exponentially.

We will now derive results for the mean, variance, and Fano factor of subcritical branching processes. Following previous results, taking expectation values of both sides of Eq. (S1) yields 〈*A*_*t*+1_〉 = *m*〈*A*_*t*_〉 + *h*. Because of stationarity 〈*A*_*t*+1_〉 = 〈*A*_*t*_〉 = 〈*A*_∞_〉 and the mean activity is given by

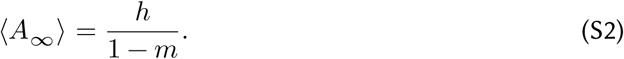

In order to derive an expression for the variance of the stationary distribution, observe that by the theorem of total variance, Var[*A*_*t*+1_] = 〈Var[*A*_*t*+1_ | *A*_*t*_]) + Var[〈*A*_*t*+1_ | *A*_*t*_〉], where 〈·〉 denotes the expected value, and *A*_*t*+1_ | *A*_*t*_ conditioning the random variable *A*_*t*+1_ on *A*_*t*_. Because *A*_*t*+1_ is the sum of independent random variables, the variances also sum: Var[*A*_*t*+1_ | *A*_*t*_] = *σ*^2^ *A*_*t*_ + *ξ*^2^. Using the previous result for 〈*A*_∞_〉 one then obtains

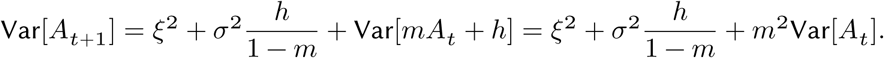

Again, in the stationary distribution Var[*A*_*t*+1_] = Var[*A*_*t*_] = Var[*A*_∞_] which yields

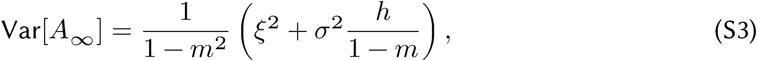

The Fano factor *F*_*A*_*t*__ = Var[*A*_*t*_]/〈*A*_*t*_〉 is easily computed from (S2) and (S3):

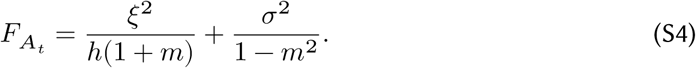

Interestingly, the mean rate, variance, and Fano factor all diverge when approaching criticality (given a constant input rate *h*): 〈*A*_∞_〉 → ∞, Var[*A*_∞_] → ∞, and *F*_*A*_*t*__ → ∞ as *m* → 1.

These results were derived without assuming any particular law for *Y* or *H*. Although the limiting behavior of BPs does not depend on it,^50–52^ fixing particular laws allows to simplify these expressions further.

We here chose Poisson distributions with means *m* and *h* for *Y* and *H* respectively: *y*_*t*, *i*_ ~ Poi(*m*) and *h*_*t*_ ~ Poi(*h*). We chose these laws for two reasons: (1) Poisson distributions allow for non-trivial offspring distributions with easy control of the branching ratio *m* by only one parameter. (2) For the brain, one might assume that each neuron is connected to *k* postsynaptic neurons, each of which is excited with probability *p*, motivating a binomial offspring distribution with mean *m* = *kp*. As in cortex *k* is typically large and *p* is typically small, the Poisson limit is a reasonable approximation. Choosing these distributions, the variance and Fano factor become

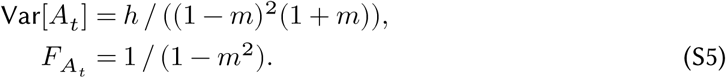

Both diverge when approaching criticality (*m* = 1).

### Supp. 2 Subsampling

A general notion of subsampling was introduced in Wilting and Priesemann [49]. The subsampled time series *a*_*t*_ is constructed from the full process *A*_*t*_ based on the three assumptions: (i) The sampling process does not interfere with itself, and does not change over time. Hence the realization of a subsample at one time does not influence the realization of a subsample at another time, and the conditional *distribution* of (*a*_*t*_|*A*_*t*_) is the same as (*a*_*t*′_|*A*_*t*′_) if *A*_*t*_ = *A*_*t*′_. However, even if *A*_*t*_ = *A*_*t*′_, the subsampled *a*_*t*_ and *a*_*t*′_ do not necessarily take the same value. (ii) The subsampling does not interfere with the evolution of *A*_*t*_, i.e. the process evolves independent of the sampling. (iii) *On average a*_*t*_ is proportional to up to a constant term, 〈*a*_*t*_ | *A*_*t*_〉 = *αA*_*t*_ + *β*.

In the spike recordings analyzed in this study, the states of a subset of neurons are observed by placing electrodes that record the activity of the same set of neurons over the entire recording. This implementation of subsampling translates to the general definition in the following manner: If *n* out of all *N* neurons are sampled, the probability to sample *a*_*t*_ active neurons out of the actual active neurons follows a hypergeometric distribution, *a*_*t*_ ~ Hyp(*N*, *n*, *A*_*t*_). As 〈*a*_*t*_ | *A*_*t*_ = *j*〉 = *jn*/*N*, this representation satisfies the mathematical definition of subsampling with *α* = *n*/*N*. Choosing this special implementation of subsampling allows to derive predictions for the Fano factor under subsampling and the spike count cross correlation. First, evaluate Var[*a*_*t*_] further in terms of *A*_*t*_:

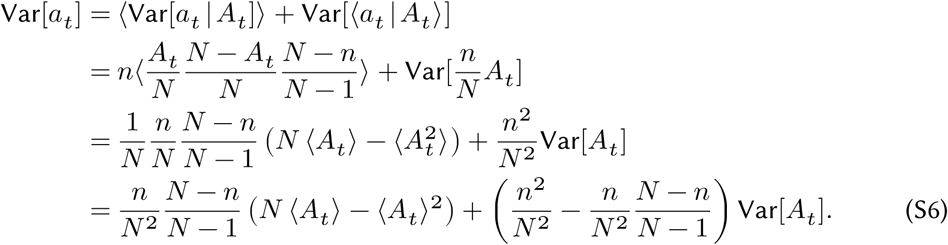

This expression precisely determines the variance Var[*a*_*t*_] under subsampling from the properties 〈*A*_*t*_〉 and Var[*A*_*t*_] of the full process, and from the parameters of subsampling *n* and *N*. We now show that the Fano factor approaches and even falls below unity under strong subsampling, regardless of the underlying dynamical state *m*. In the limit of strong subsampling (*n* ≪ *N*) Eq. (S6) yields:

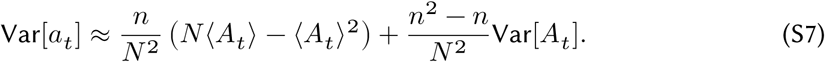

Hence the subsampled Fano factor is given by

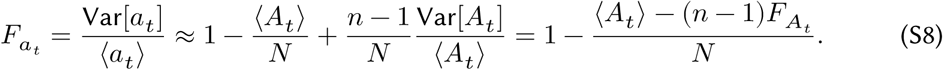

Interestingly, when sampling a single unit (*n* = 1) the Fano factor of that unit becomes completely independent of the Fano factor of the full process:

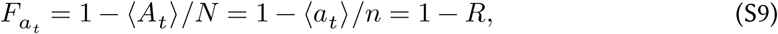

where *R* = 〈*a*_*t*_〉/*n* is the mean rate of a single unit.

Based on this implementation of subsampling, we derived analytical results for the crosscorrelation between the activity of two units on the time scale of one time step. The pair of units is here represented by two independent samplings *a*_*t*_ and *ã*(*t*) of a BP *A*_*t*_ with *n* = 1, i. e. each represents one single unit. Because both samplings are drawn from identical distributions, their variances are identical and hence the correlation coefficient is given by *r*_sc_ = Cov(*a*_*t*_, *ã*(*t*))/Var[*a*_*t*_]. Employing again the law of total expectation and using the independence of the two samplings, this can be evaluated:

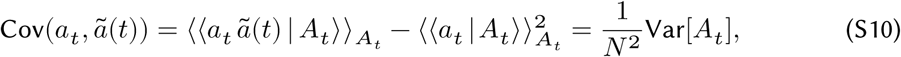

with the first inner expectation being taken over the joint distribution of *a*_*t*_ and *á*(*t*). Using Eq. (S7), one easily obtains

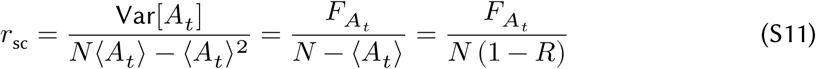

with the mean single unit rate *R* = 〈*A*_*t*_〉/*N*. For subcritical systems, the Fano factor *F*_*A*_*t*__ is much smaller than *N*, and the rate is typically much smaller than 1. Therefore, the crosscorrelation between single units is typically very small.

**Figure S1:**
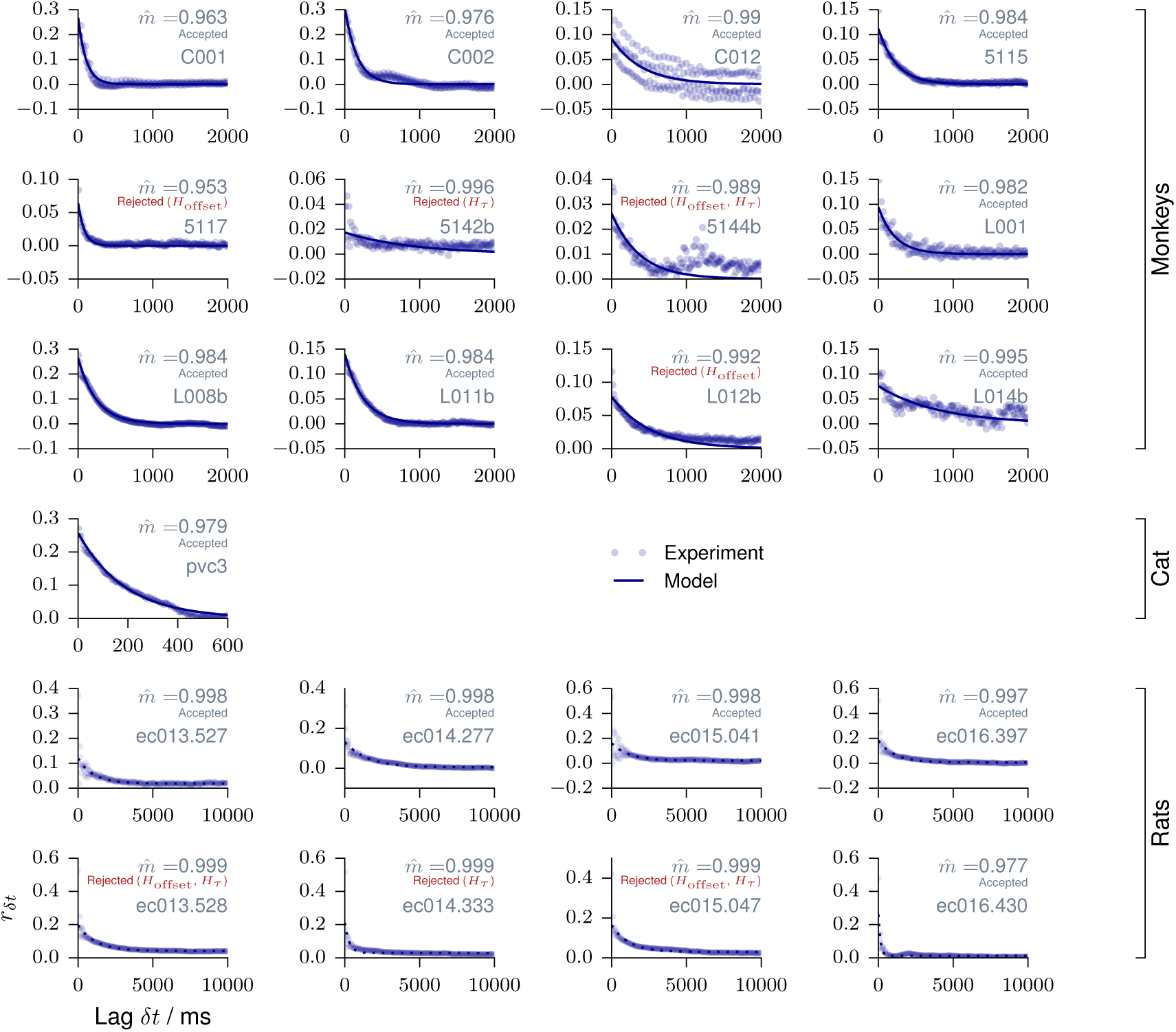
MR estimation for individual recording sessions. Reproduced from^49^. MR estimation is shown for every individual animal. The consistency checks are detailed in^49^. Data from monkey were recorded in prefrontal cortex during an working memory task. The third panel shows a oscillation of *r*_*k*_ with a frequency of 50 Hz, corresponding to measurement corruption due to power supply frequency. Data from anesthetized cat were recorded in primary visual cortex. Data from rat were recorded in hippocampus during a foraging task. In addition to a slow exponential decay, the slopes *r*_*k*_ show the *ϑ*-oscillations of 6 - 10 Hz present in hippocampus.

**Figure S2:**
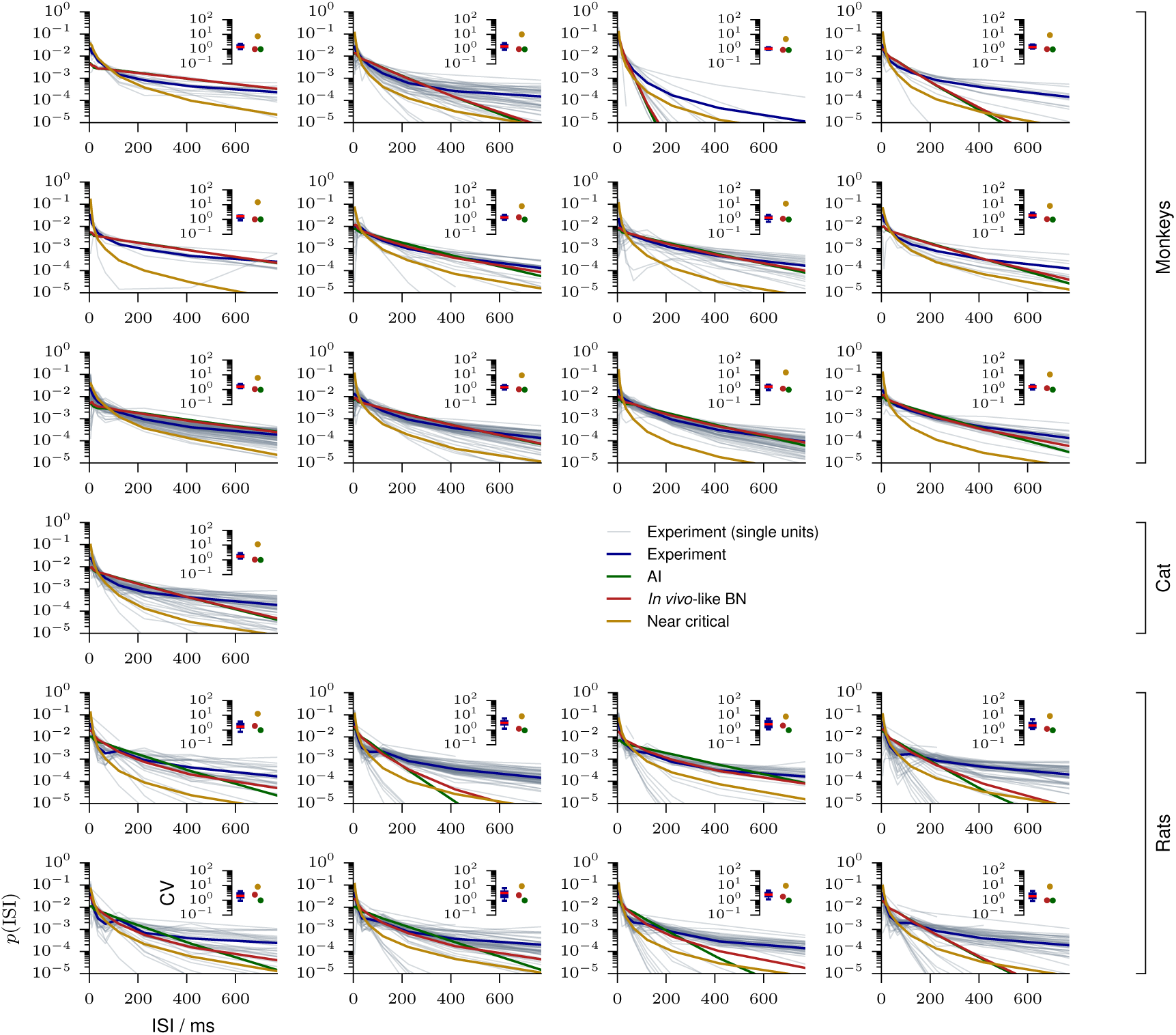
Interspike interval distribution for individual recording sessions. Interspike interval (ISI) distributions are shown for individual units of each recording (gray), for the average over units of each recording (blue), as well as for the matched models, either AI (green), *in vivo*-like (red), or near critical (yellow). The insets show the corresponding coefficients of variation (CV). For every experiment AI and *in vivo*-like models are virtually indistinguishable by the ISI distributions.

**Figure S3:**
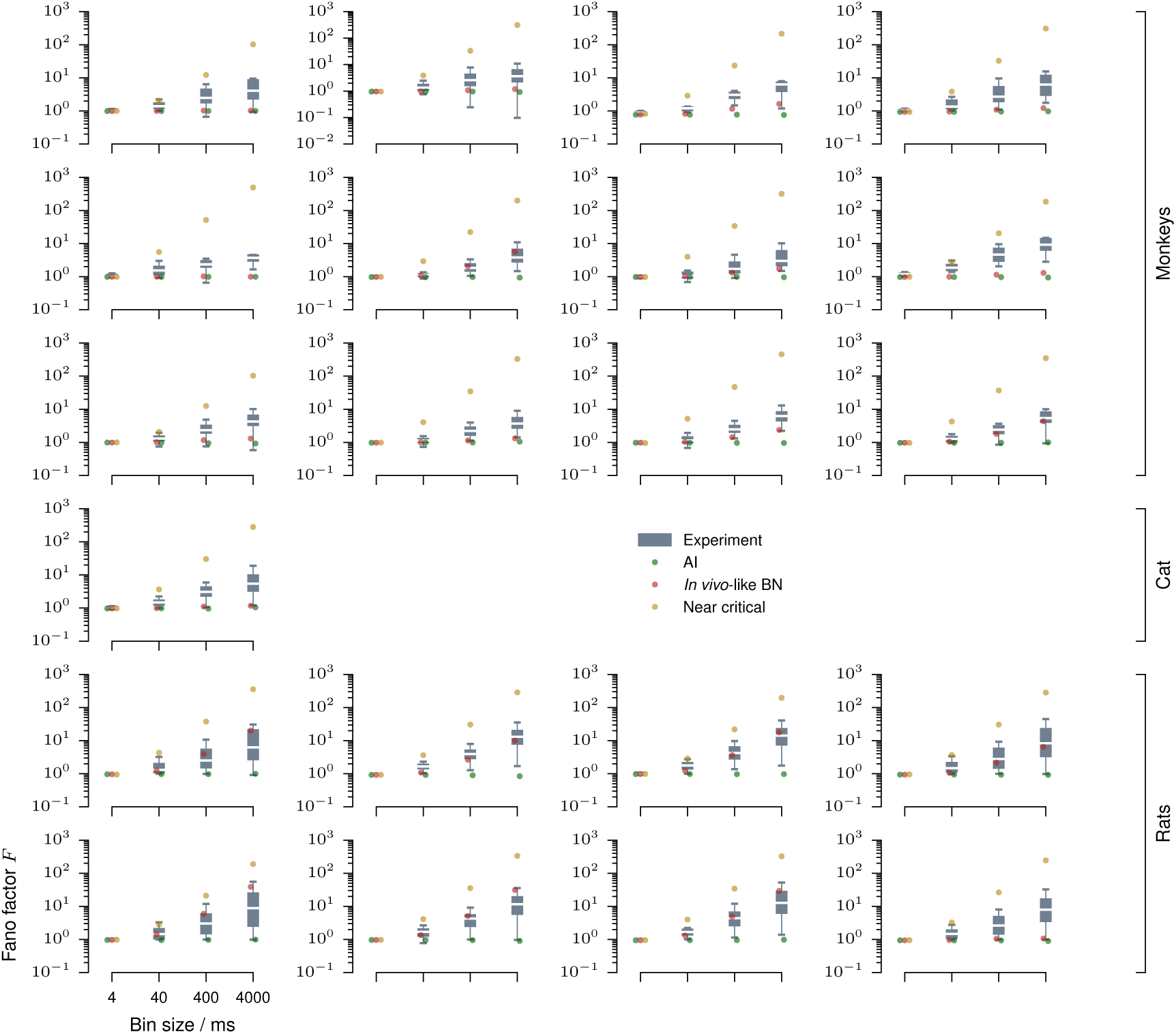
Fano factors for individual recording sessions. Fano factors are shown for individual single or multi units of every recording (gray boxplots, median / 25% - 75%, 2.5% - 97.5%), as well as for the matched models, either AI (green), *in vivo*-like (red), or near critical (yellow).

**Figure S4:**
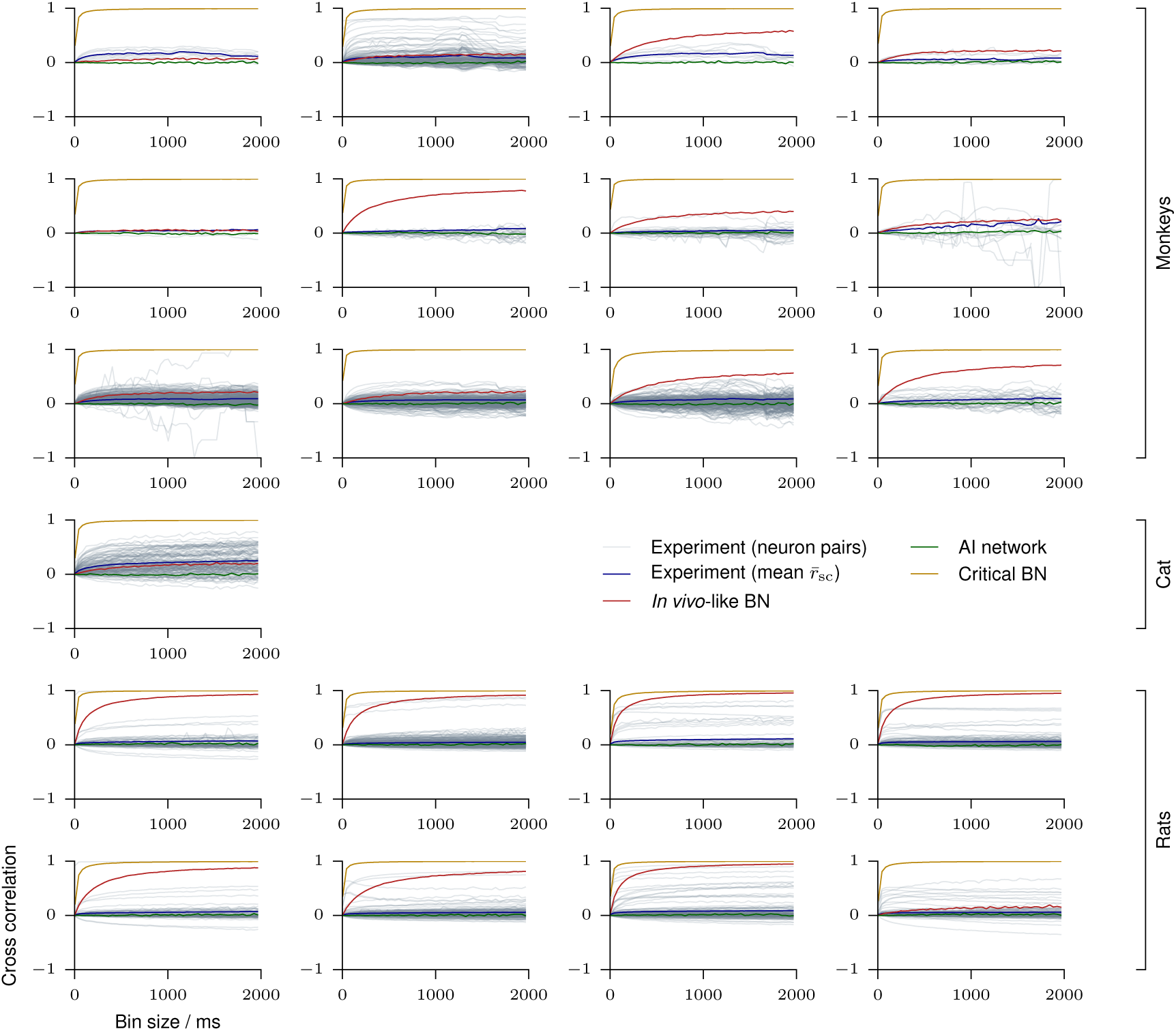
Cross correlations for individual recording sessions. Spike count cross correlations (*r*_sc_) are shown for every neuron pair (gray) and the ensemble average (blue) of each recording, for bin sizes from 1 ms to 2s. Cross correlations are also shown for the matched models, either AI (green), *in vivo*-like (red), or near critical (yellow).

**Figure S5:**
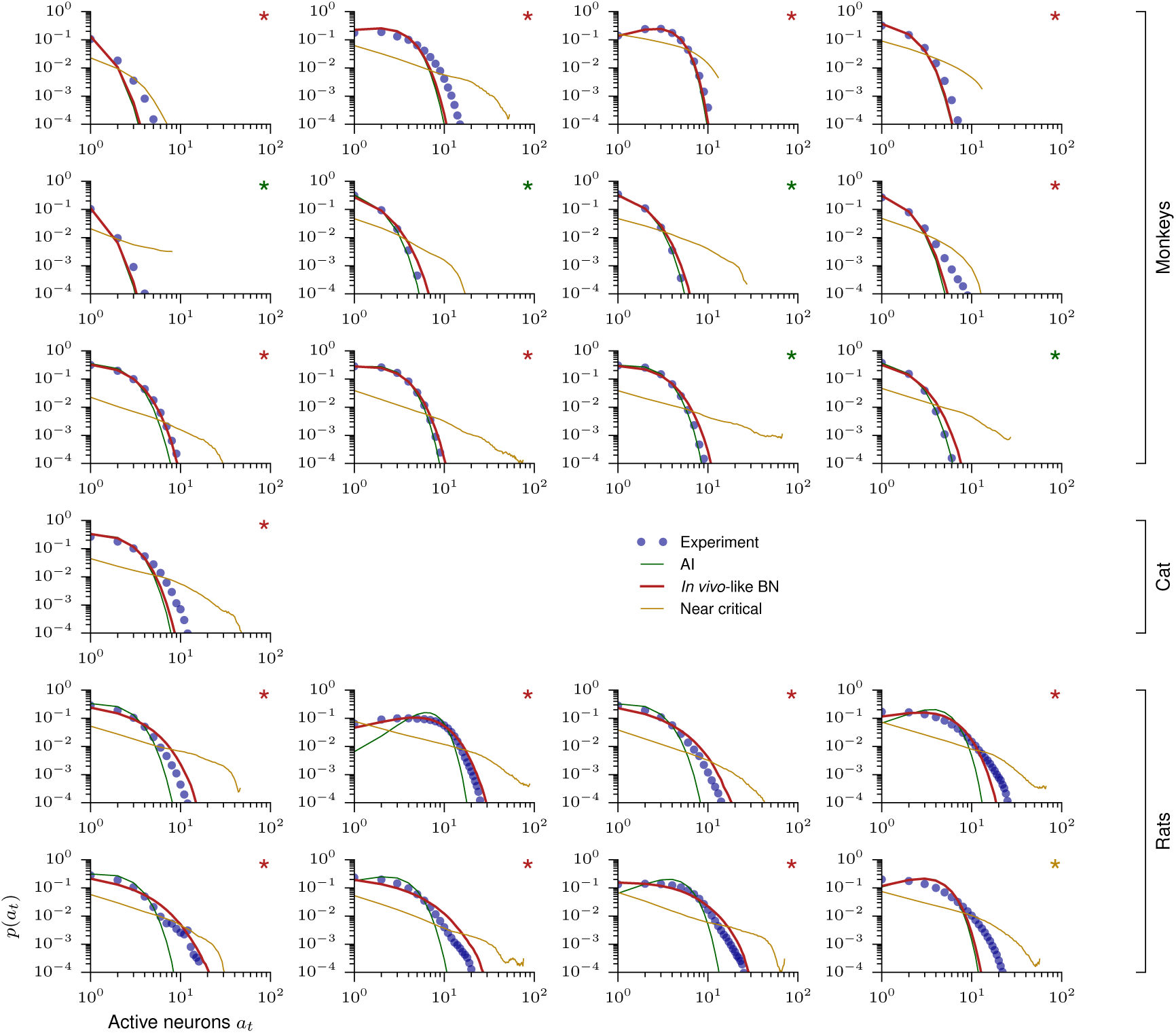
Activity distributions (4 ms bin size). Activity distributions are shown for every recording for a bin size of 4 ms (blue). Activity distributions for the matched models, either AI (green), *in vivo*-like (red), or near critical (yellow) are also shown. The color of the asterisk indicates which of the three models yielded the highest likelihood for the data following^64^.

**Figure S6:**
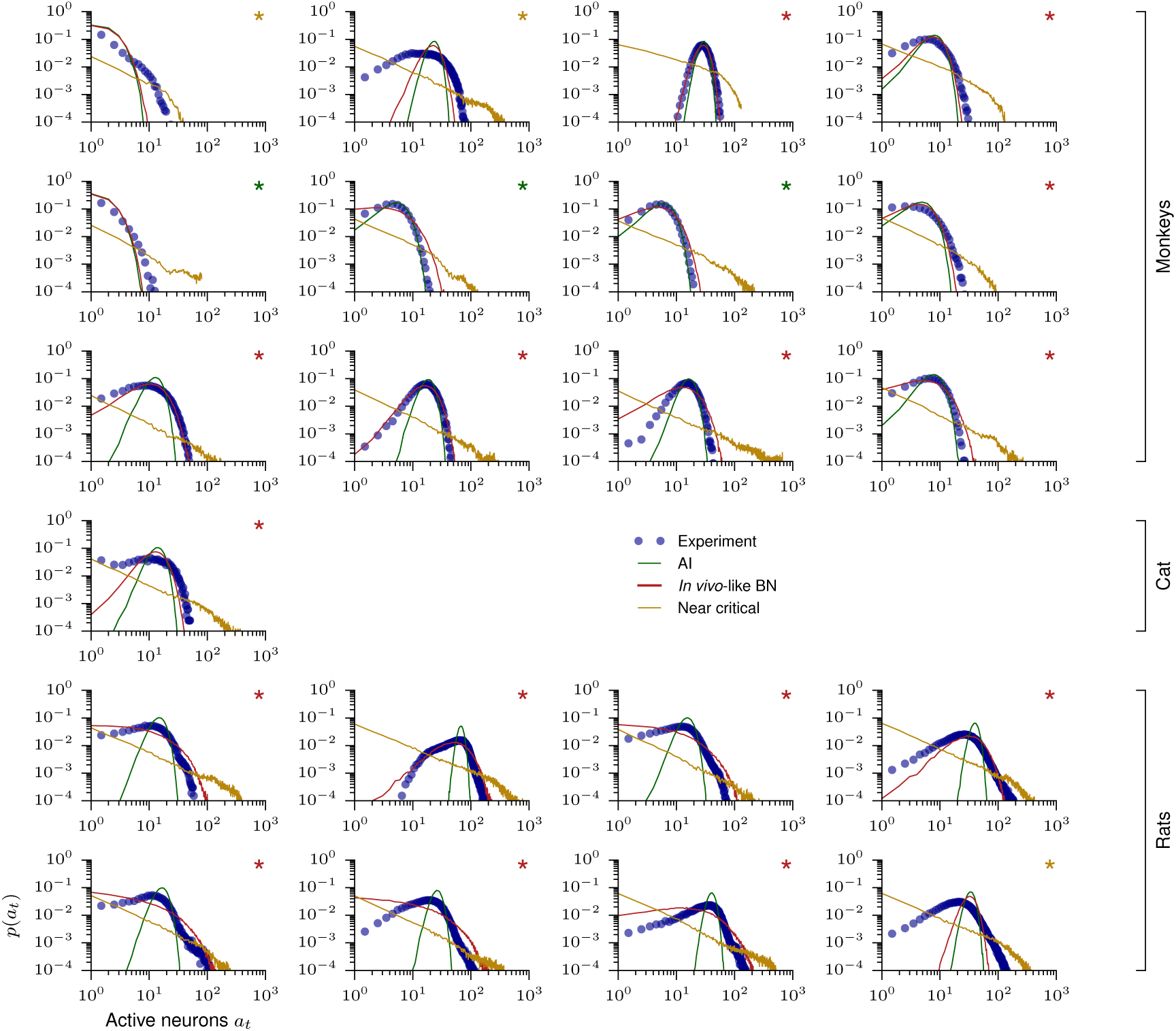
Activity distributions (40 ms bin size). Activity distributions are shown for every recording, for a bin size of 40 ms (blue). Activity distributions for the matched models, either AI (green), *in vivo*-like (red), or near critical (yellow) are also shown. The color of the asterisk indicates which of the three models yielded the highest likelihood for the data following^64^.

**Figure S7:**
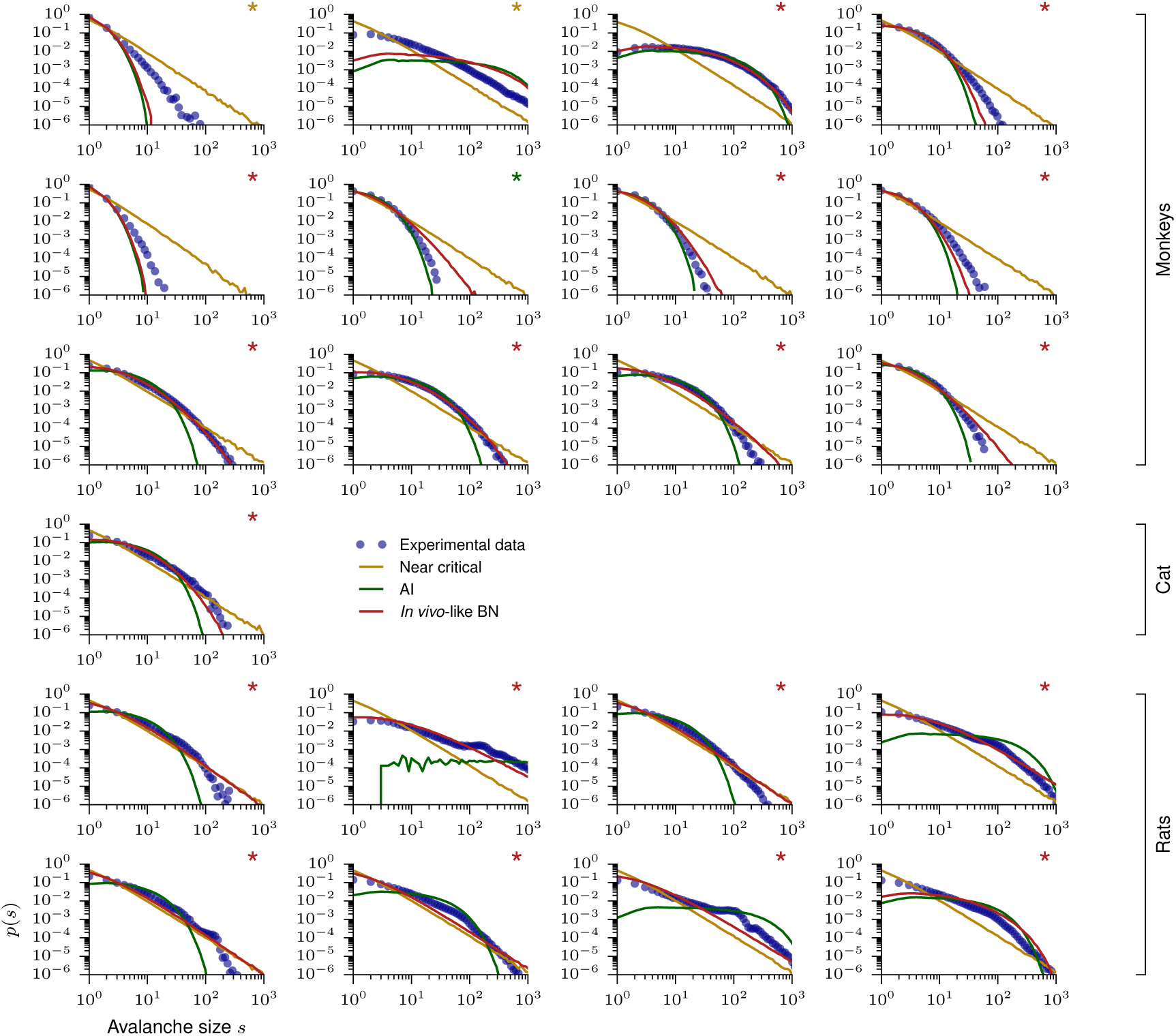
Avalanche size distribution for individual recording sessions. Avalanche size distributions are shown for every recording (blue) and for matched models, either AI (green), *in vivo*-like (red), or near critical (yellow). The color of the asterisk indicates which of the three models yielded the highest likelihood for the data following^64^.

**Figure S8:**
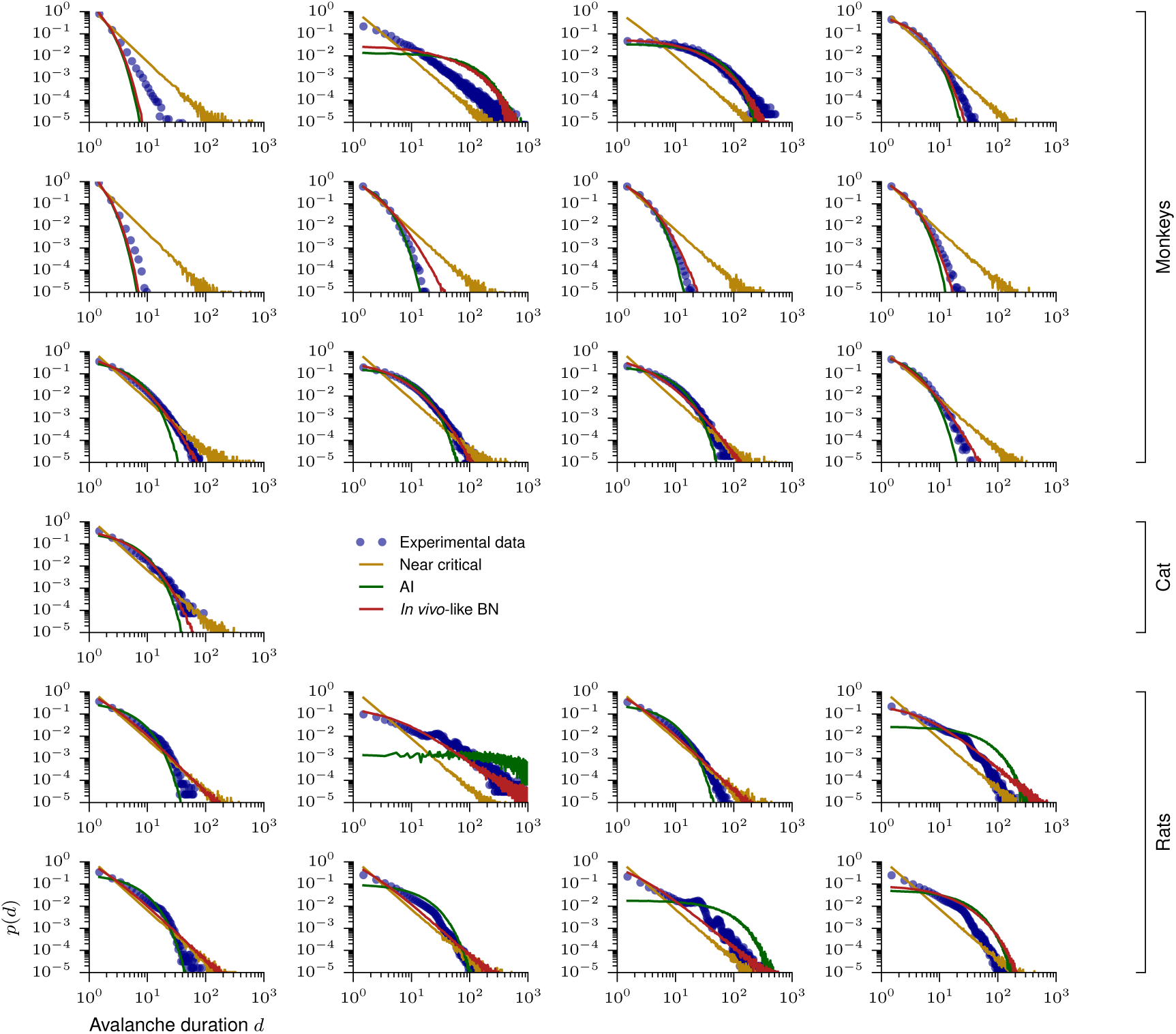
Avalanche duration distribution for individual recording sessions. Avalanche duration distributions are shown for every recording (blue) and for matched models, either AI (green), *in vivo*-like (red), or near critical (yellow).

**Figure S9:**
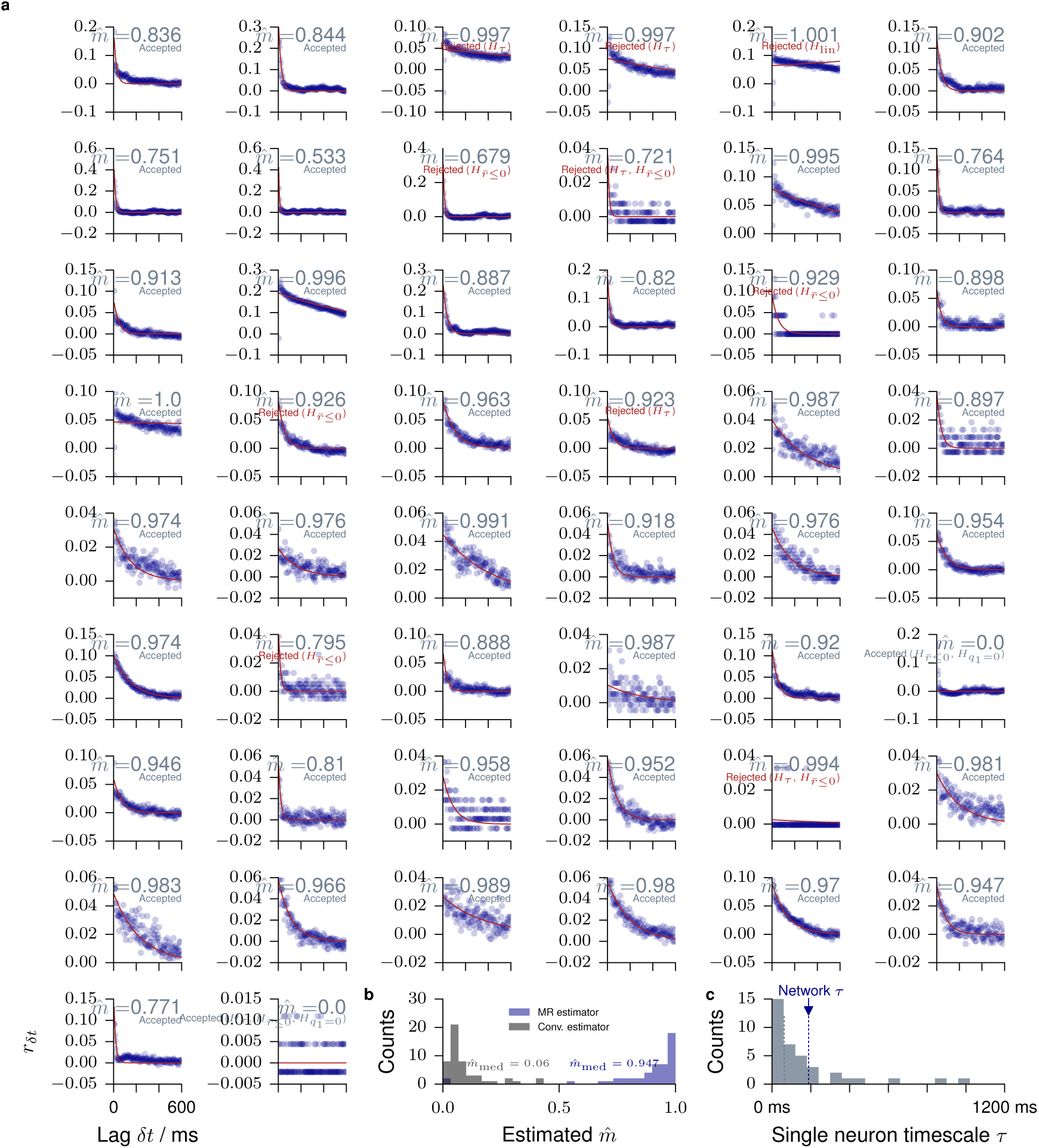
MR estimation from single neuron activity (cat). Modified from^49^. MR estimation is used to estimate *m*̂ from the activity *a*_*t*_ of a single units in cat visual cortex. **a**. Each panel shows MR estimation for one of the 50 recorded units. Autocorrelations decay rapidly in some units, but long-term correlations are present in the activity of most units. The consistency checks are detailed in^49^. **b**. Histogram of the single unit branching ratios *m*̂, inferred with the conventional estimator and using MR estimation. The difference between these estimates demonstrates the subsampling bias of the conventional estimator, and how it is overcome by MR estimation. **c**. Histogram of single unit timescales with their median (gray dotted line) and the timescale of the dynamics of the whole network (blue dotted line).

**Figure S10:**
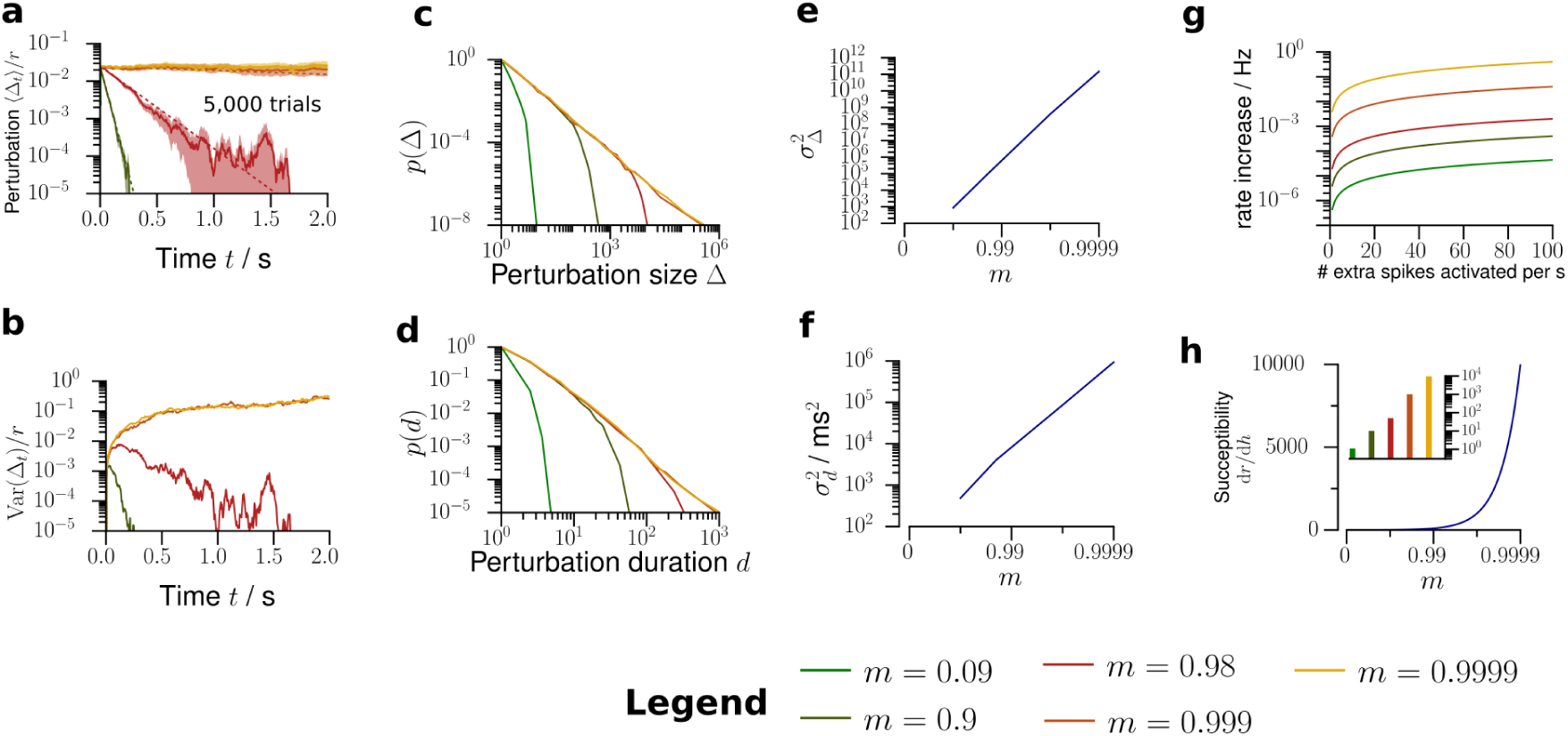
Further predictions about network activity. **a**. The model predicts that the perturbation decays exponentially with decay time *τ* = –Δ*t*/log *m*. **b** The variance across trials of the perturbed firing rate has a maximum, whose position depends on *m*. **c**. Depending on *m*, the model predicts the distributions for the total number of extra spikes *Δ* generated by the network following a single extra spike. **d**. Likewise, the model predicts distributions of the duration *d* of these perturbations. **e**. Variance of the total perturbation size as a function of *m*. **f**. Variance of the total perturbation duration as a function of *m*. **g**. Increase of the network firing rate as a function of the rate of extra neuron activations for different *m*. **h**. Amplification (susceptibility) d*r*/d*h* of the network as a function of the branching ratio *m*.

